# Compositional Shifts and Co-occurrence Patterns of Topsoil Bacteria and Micro-Eukaryotes Across a Permafrost Thaw Gradient in Alpine Meadows of the Qilian Mountains, China

**DOI:** 10.1101/2024.05.14.594190

**Authors:** Zhu Wang, Yang Liu, Fang Wang

## Abstract

Soil microorganisms play a pivotal role in the biogeochemical cycles of alpine meadow ecosystems, especially in the context of permafrost thaw. However, the mechanisms driving microbial community responses to environmental changes, such as variations in active layer thickness (ALT) of permafrost, remain poorly understood. This study utilized next-generation sequencing to explore the composition and co-occurrence patterns of soil microbial communities, focusing on bacteria and micro-eukaryotes along a permafrost thaw gradient. Results showed a decline in bacterial alpha diversity with increasing permafrost thaw, while micro-eukaryotic diversity exhibited an opposite trend. Although changes in microbial community composition were observed in permafrost and seasonally frozen soils, these shifts were not statistically significant. Bacterial communities exhibited greater differentiation between frozen and seasonally frozen soils, a pattern not mirrored in eukaryotic communities. LEfSe analysis revealed a higher number of potential biomarkers in bacterial communities compared to micro-eukaryotes. Bacterial co-occurrence networks were more complex, with more nodes, edges, and positive linkages than those of micro-eukaryotes. Key factors such as soil texture, ALT, and bulk density significantly influenced bacterial community structures, particularly affecting the relative abundances of the Acidobacteria, Proteobacteria, and Actinobacteria phyla. In contrast, fungal communities (e.g., *Nucletmycea*, *Rhizaria*, *Chloroplastida*, and *Discosea* groups) were more affected by electrical conductivity, vegetation coverage, and ALT. This study highlights the distinct responses of soil bacteria and micro-eukaryotes to permafrost thaw, offering insights into microbial community stability under global climate change.

**Importance:** This study sheds light on how permafrost thaw affects microbial life in the soil, with broader implications for understanding climate change impacts. As permafrost degrades, it alters the types and numbers of microbes in the soil. These microbes play essential roles in environmental processes, such as nutrient cycling and greenhouse gas emissions. By observing shifts from bacteria-dominated to fungi-dominated communities as permafrost thaws, the study highlights potential changes in these processes. Importantly, this research suggests that the stability of microbial networks decreases with permafrost degradation, potentially disrupting the delicate balance of these ecosystems. The findings not only deepen our understanding of microbial responses to changing climates but also support the development of strategies to monitor and potentially mitigate the effects of climate change on fragile high-altitude ecosystems.

## Introduction

Permafrost, covering approximately 18% of the Northern Hemisphere (1, 2), plays a crucial role in global carbon cycling due to its storage of an estimated 15% of the world’s soil carbon (3). Recent measurements showed that global permafrost has warmed significantly by 0.29L (4), which has led to its accelerated degradation. This degradation is evidenced by reductions in extent and thickness, as well as increases in the depth of the seasonally thawed active layer (5, 6, 7). These changes disrupt regional thermo-hydrological processes and release stored soil organic carbon, thus affecting ecosystem functions.

Microorganisms in permafrost are pivotal to the decomposition and mineralization of organic matters, significantly influencing biogeochemical cycles (8, 9). Microbial communities respond swiftly to permafrost thaw (10), transforming ecosystems from carbon sinks to carbon sources by producing greenhouse gases, such as carbon dioxide and methane, through enhanced decomposition activities (11). This shift is accompanied by increased transcriptional activity and methane production (12). As global warming progresses, permafrost degradation modifies heat and nutrient dynamics within the active layer, affecting microbial community structure, phylogeny, functional genes, and metabolic pathways (13). The thawing process makes previously locked soil organic carbon bioavailable, intensifying microbial metabolism and accelerating the breakdown of organic matter, thereby augmenting greenhouse gas emissions (14). Understanding these microbial dynamics is essential for predicting how permafrost ecosystems will respond to climate change.

The Qinghai-Tibet Plateau, often referred to as the “Earth’s Third Pole” and “Asia’s Water Tower”, hosts the most extensive permafrost distribution in low and middle latitudes. The permafrost in this region is thawing more rapidly than its high-latitude counterparts in North America, making it particularly vulnerable to climate change (5). At the beginning of the 21st century, permafrost on the Qinghai-Tibet Plateau covered approximately 1.5 million km^2^, accounting for 69.8% of China’s total permafrost area (15, 16, 17). Recent studies indicate a significant reduction in permafrost coverage, declining by 66,000 km^2^ per decade from 1980 to 2010 (18). The Qilian Mountains, situated on the northern edge of the plateau, are highly sensitive to climatic fluctuations, making them an important area for studying permafrost ecosystem responses to global warming. Previous studies on soil microbial dynamics in permafrost regions have primarily focused on environmental gradients, such as elevation (19, 20) and vegetation types (21, 22). These studies have linked key soil properties, such as moisture, pH, and organic carbon, to microbial community shifts (23, 24). However, vegetation type and elevation, as composite environmental factors, encompass various interrelated elements, and their impacts on microbial communities generally manifesting as long-term, cumulative effects. For example, vegetation type influences soil organic matter input, as well as soil temperature and moisture (25), while elevation indirectly affects soil conditions by altering climate parameters (26). In contrast, the active layer thickness (ALT) of permafrost directly affects soil physical and chemical properties, underscoring its unique significance in microbial community composition and function (27). Studying the impact of ALT on microbial communities provides insights into how thawing alters soil conditions, affecting microbial diversity and activity (28, 29). This understanding is crucial for predicting how changing permafrost conditions will influence nutrient cycling, carbon and nitrogen release, and overall ecosystem functioning in alpine meadows.

Our study addresses the gap in understanding how microbial communities respond to permafrost thaw by examining compositional shifts in bacterial and micro-eukaryotic communities along a gradient of active layer thickness (ALT) in the Qilian Mountains. By employing a space-for-time substitution approach, we investigated how shifts in microbial communities correlate with changes in soil physical and chemical properties across different stages of permafrost thaw. The specific objectives of our research are to: (i) Evaluate the alpha diversity and compositional shifts of bacterial and micro-eukaryotic communities in permafrost and seasonally frozen soils; (ii) Analyze co-occurrence patterns and determine the relative importance of various bacterial and micro-eukaryotic taxa in shaping microbial networks within permafrost ecosystems; (iii) Investigate the key environmental factors influencing microbial diversity and community structure across different stages of permafrost thaw. By examining these aspects, our research aims to deepen the understanding of how soil microbial communities adapt to changing permafrost conditions and contribute to biogeochemical cycling in alpine meadow ecosystems. This insight is crucial for predicting and managing the impacts of climate change on permafrost regions globally.

## 1 Materials and methods

### 1.1 Study sites and soil sample collection

The Heihe River basin, located on the northeastern edge of the Qilian Mountains, is the second-largest inland river system in China and is currently experiencing significant climatic warming (30, 31). In August 2019, a comprehensive soil sampling campaign was conducted at eight strategic sites along the upper reaches of the Heihe River, with geographic coordinates ranging from 98°40′E to 99°20′E and from 38°30′N to 38°60′N (Fig. 1). These sites are situated at altitudes between 3400 to 4700 meters above sea level. The mean annual precipitation (MAP) and mean annual temperature (MAT) in this region range from 200 to 500 mm (32) and from –2.5 to –0.3 L (33), respectively. The permafrost in the Qilian Mountains exhibits substantial environmental variability, with ground temperature being a key indicator of permafrost degradation (34). Based on established classifications of permafrost types determined by annual mean ground temperature (35, 36), soil samples were classified as permafrost (sites N1, N2, N3, N4, and N5) or seasonally frozen soils (sites L1, L2, and L3). This classification enabled targeted investigations of microbial community responses to different degrees of permafrost degradation, with additional data on active layer thickness sourced from the Chinese National Tibetan Plateau Data Center (http://data.tpdc.ac.cn, supported by 35).

**Fig. 1.**
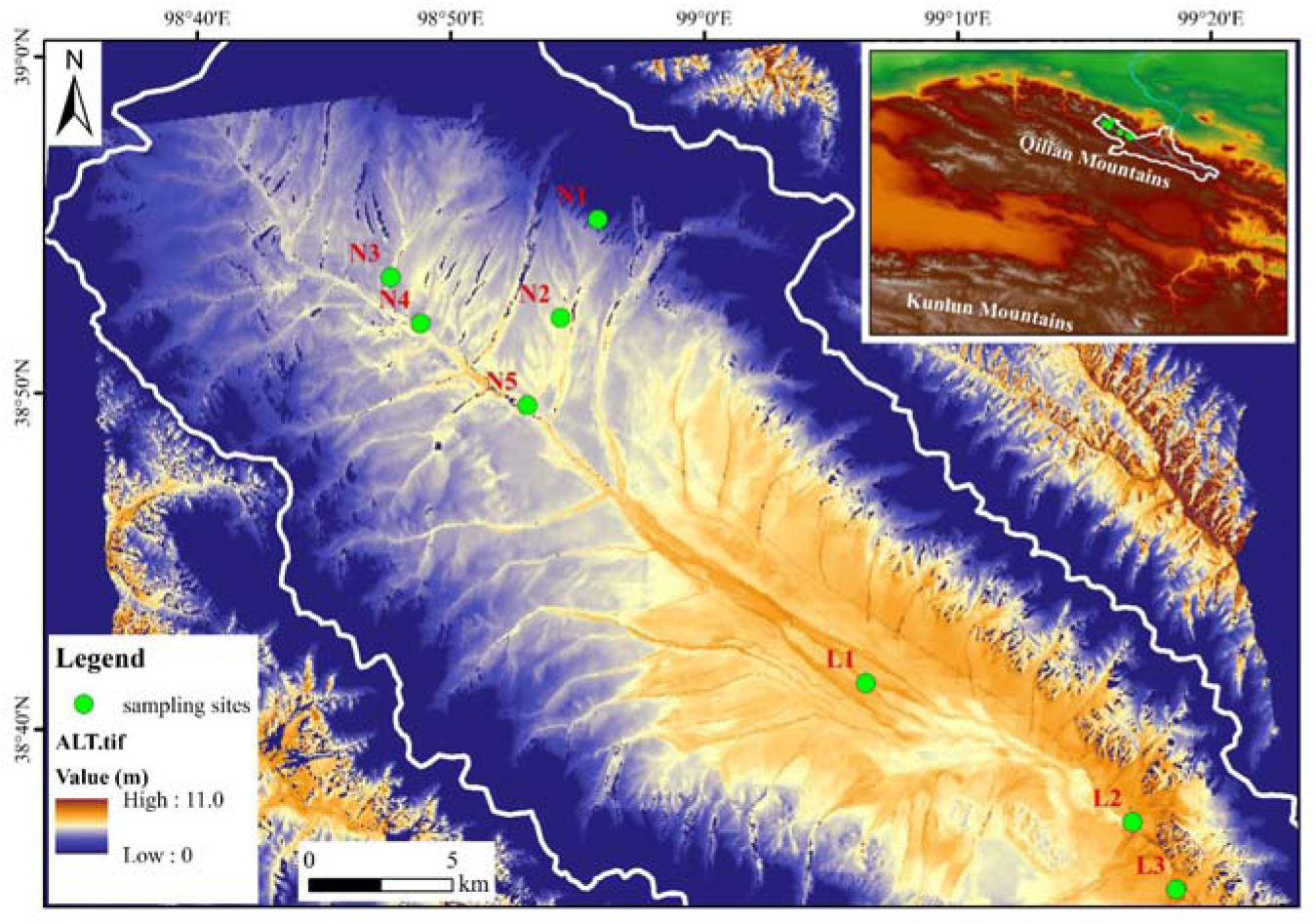
Sampling sites with locations spanning permafrost soils (N1-N5) and seasonally frozen soils (L1-L3) in the upstream region of the Heihe River.

Vegetation along the river’s course varied across the study sites. Near the river’s origin (N1, N2, and N3), alpine meadows were dominated by grasses, such as *Polygonum viviparum*, *Lagotis brachystachya*, and *Rhodiola algida*, are prevalent. Further downstream, the vegetation shifted to swamp meadows (N4, N5) and steppe meadows (L1, L2, and L3), dominated by *Kobresia capillifolia* and *Carex moorcroftii*, respectively. Detailed information on permafrost parameters and vegetation at each site was provided in Table S1.

A total of 24 soil samples were systematically extracted from three distinct layers (0-20 cm, 20-40 cm, and 40-50cm) at each site. Samples were collected from five random points within a 5 m × 5 m plot at each site and pooled to create composite samples representing each depth interval. Soil particle size analysis was performed at 10 cm intervals across the 0-50 cm soil profile. All samples were thoroughly mixed and sieved to remove stones and plant debris before analysis. Samples were sealed in sterilized plastic bags, kept cool with ice, and transported to the laboratory within 48 hours. Once in the laboratory, each sample was divided: one part was stored at 4 L for immediate physical and chemical analyses, while the other was cryogenically preserved at –80 L in sterilized polyethylene tubes for DNA extraction and PCR amplification.

### 1.2 Analysis of soil physicochemical properties

A detailed assessment of soil physicochemical properties was conducted, including measurements of both physical (e.g., bulk density, moisture content, temperature, electrical conductivity, particle size distribution) and chemical parameters (e.g., pH, total organic carbon, total nitrogen, NO_3_^-^-N, and NH_4_^+^-N). In situ measurements of soil moisture, temperature, and electrical conductivity were performed at each sampling depth using a portable W.E.T. Soil Moisture Meter (DELTA Inc., UK). Bulk density was determined using the core cutter method, where soil samples were excavated using a cylindrical core and oven-dried at 90 L for 24 hours. Particle size composition was assessed through a rapid sieving procedure (37).

Chemical analyses involved mixing soil samples with water at a weight-to-volume ratio of 1:5 for pH measurement, using a Leici pH electrode (Shanghai, China). Total organic carbon (TOC) and total nitrogen (TN) were quantified using dichromate digestion (38) and the Kjeldahl digestion method (39), respectively. Soil nitrate nitrogen (NO_3_ ^-^-N) and ammonium nitrogen (NH_4_ ^+^-N) concentrations were measured using KCl extraction, followed by indophenol blue colorimetry and the phenol disulfonic acid method, respectively (40). All measurements were performed in triplicate for each sample to ensure reliability and reproducibility. Average values from these measurements are provided in Table S1 for further interpretation and correlation with microbial community analyses.

### 1.3 Soil DNA extraction and PCR amplification

Total DNA was extracted from 30-50 ng of fresh soil using the MetaVx^TM^ Library Preparation Kit (GENEWIZ, Inc., South Plainfield, NJ, USA) according to the manufacturer’s protocol. DNA concentration was quantified using a Qubit2.0 Fluorometer (Invitrogen, Carlsbad, CA, USA), and its integrity was confirmed via electrophoresis on a 1.5% agarose gel in 0.5×TAE buffer.

For bacterial and archaeal 16S rDNA amplification, primers targeting the V3-V4 hypervariable regions were used: forward primer “CCTACGGRRBGCASCAGKVRVGAAT” and reverse primer “GGACTACNVGGGTWTCTAATCC” (41). For micro-eukaryotic 18S rDNA amplification, the V7-V8 hypervariable regions were amplified using the forward primer “CGWTAACGAACGAG” and the reverse primer “AICCATTCAATCGG” (42). These primers also contained adaptor sequences for high-complexity library amplification and sequencing on the Illumina MiSeq platform.

PCR amplifications were performed in triplicate in 25 μL reaction mixtures containing 2.5 μL of TransStart Buffer, 2 μL of dNTPs, 1 μL of each primer, and 20 ng of template DNA. The thermal profile for amplifying 16S rRNA fragments consisted of an initial denaturation at 94 L for 3 minutes, followed by 24 cycles of denaturation at 94 L for 5 seconds, annealing at 57 L for 90 seconds, and elongation at 72 L for 10 seconds, with a final elongation step at 72 L for 5 minutes. For 18S rRNA, the protocol included an initial denaturation at 94 L for 5 minutes, followed by 25 cycles of denaturation at 94 L for 30 seconds, annealing at 57 L for 30 seconds, and elongation at 72 L for 30 seconds, ending with a final elongation at 72 L for 5 minutes. Both PCR products were subsequently purified and quantified to prepare composite DNA samples for library construction, which were sequenced on an Illumina MiSeq platform (Illumina, Inc., San Diego, CA, USA) at GENEWIZ, Inc. (Suzhou, China).

### 1.4 High-throughput sequencing and data processing

High-throughput sequencing was conducted on an Illumina MiSeq platform using the PE250 paired-end configuration. Image analysis and base calling were managed by the MiSeq Control Software (MCS). Raw sequence data were initially processed using the Quantitative Insights into Microbial Ecology (QIIME) software package (version 1.9.1) (43). Sequences were filtered to retain only those longer than 200bp and devoid of ambiguous bases (“N”). The high-quality sequences were then clustered into operational taxonomic units (OTUs) at 97% sequence similarity using the VSEARCH algorithm (version 1.9.6). Taxonomic assignment of each OTU for both 16S rDNA and 18S rDNA sequences was performed using the Ribosomal Database Project (RDP) classifier (version 2.2; http://sourceforge.net/project/rdp-classifier/), with a confidence threshold set at 0.8 (44). The RDP classifier utilized the Silva 138 database (Release 138; http://www.arb-silva.de), enabling taxonomic predictions down to the species level for reliable annotation of microbial community structure. Raw sequencing reads were uploaded to the NCBI Sequence Read Archive (SRA) database. Accession numbers for 16S rRNA are SRR30963207, SRR30963205, SRR30963206, SRR30963208, SRR30963204, SRR30963201, SRR30963202, and SRR30963203. For 18S rRNA data, the accession numbers are SRR30961751, SRR30961750, SRR30961748, SRR30961749, SRR30961747, SRR30961744, SRR30961745, and SRR30961746.

### 1.5 Statistical analysis

The *t*-test was used to detect significant differences in soil physicochemical properties and microbial community relative abundances between permafrost soils (group N) and seasonally frozen soils (group L), with significance set at *P* <0.05. Changes in relative abundances of bacterial and micro-eukaryotic communities across the permafrost thaw gradient were analyzed using generalized linear models in SPSS (version 25.0, IBM Corp., Chicago, IL, USA). Microbial alpha-diversity indices, including Richness, Chao1, and Shannon, were computed from rarefied OTU matrices using the “phyloseq” package in R (v.3.6.2) (45). Boxplots illustrating these indices were generated in R for visualizing diversity patterns. The structure of bacterial and micro-eukaryotic communities was assessed using ecological distance measures and multivariate statistical methods in R. Community comparisons were performed using UniFrac distances (46). Beta-diversity between sampling sites was analyzed through nonmetric multidimensional scaling (NMDS) and Analysis of Similarities (ANOSIM) based on Bray-Curtis distances to reflect variations in OTU composition. This analysis was used to compare microbial community structures between different soil conditions, with beta-diversity capturing how dissimilar or similar the microbial communities were across permafrost and seasonally frozen soils. Linear Discriminant Analysis Effect Size (LEfSe) was used to identify differentially abundant microbial biomarkers between permafrost and seasonally frozen soils, with the Kruskal-Wallis test applied (LDA score > 2.0, *P* < 0.05). Co-occurrence networks for bacterial and micro-eukaryotic communities were constructed using the IDENAP process (http://mem.rcees.ac.cn:8081), elucidating intra-domain associations (47). OTUs for network analysis were selected based on relative abundance (>0.01%) and occurrence frequency (>50% of samples), focusing on ecologically significant taxa. A correlation coefficient matrix with a threshold cutoff of 0.94 (*P* < 0.05), as determined by a cutoff plot, ensured robust network relationships. Networks were visualized using Gephi (version 0.9.6) (48). The relationship between soil properties and microbial community compositions was examined using Mantel tests, conducted with the “ggcor” and “vegan” packages in R. Redundancy Analysis (RDA) was performed in CANOCO 4.5 for Windows (Biometris, Wageningen, Netherlands) to identify environmental drivers influencing microbial community differentiation at the phylum level, with significance assessed using a Monte Carlo permutation test (999 permutations).

## 2 Results

### 2.1 Variations in the soil physicochemical properties

The variations in soil physicochemical properties across permafrost soils (sites N1-N5) and seasonally frozen soils (sites L1-L3) at three depth intervals were presented in Figure 2. Along the soil profile, bulk density (BD) and electrical conductivity (EC) showed an increasing trend, particularly in the permafrost soils, although these changes were not statistically significant (*P* > 0.05). Soil temperature (ST) consistently decreased with depth, following a predictable gradient. Soil moisture (SM) was higher in the surface layers, with an unusual inversion observed at sites L2 and L3. The pH values ranged from 7.80 to 8.55, indicating uniformly alkaline conditions across all samples. Total organic carbon (TOC) and total nitrogen (TN) varied between 2.0 to 18.5 g/kg and 1.0 to 2.5 g/kg, respectively, with TOC generally decreasing with depth. TN, nitrate (NN), and ammonia (AN) levels did not exhibit consistent trends across the soil profile, suggesting localized influences on nitrogen dynamics at individual sites.

**Fig. 2.**
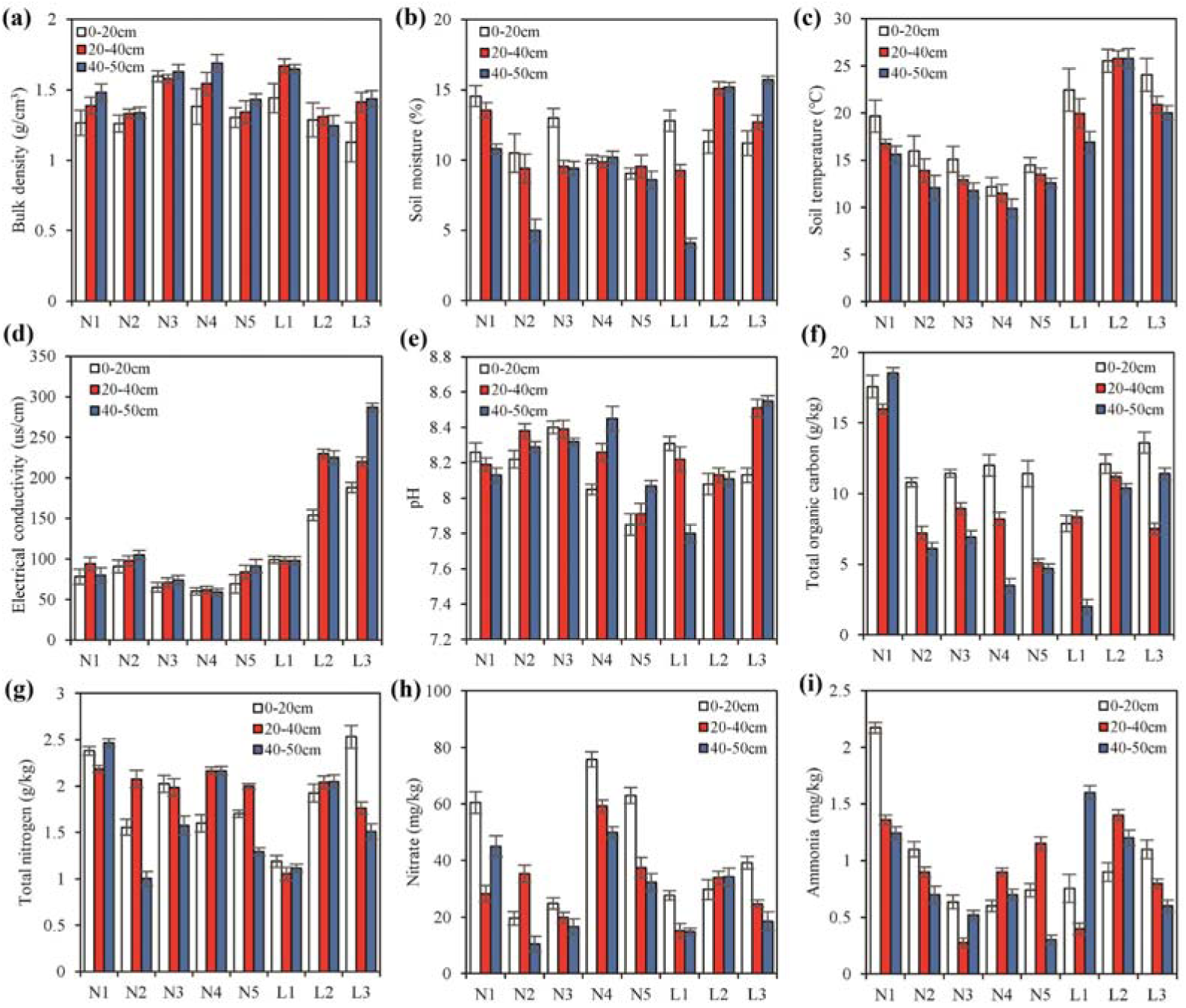
Soil properties of permafrost soils (N1-N5) and seasonally frozen soils (L1-L3) under different layers (white: 0-20 cm, red: 20-40 cm, blue: 40-50 cm) at sampling sites: (a) Bulk density, (b) Soil moisture, (c) Soil temperature, (d) Electrical conductivity, (e) pH, (f) Total organic carbon, (g) Total nitrogen, (h) Nitrate, and (i) Ammonia. All data are presented as “mean ± standard deviation” calculated from triplicated samples.

Results from the *t*-test (Fig. S1) indicated that only soil temperature and electrical conductivity differed significantly between permafrost and seasonally frozen soils, with both parameters showing higher mean values in seasonally frozen soils. No statistically significant differences were observed for the other soil physiochemical properties. However, permafrost soils generally had higher average concentrations of TOC, TN, and NN compared to seasonally frozen soils, whereas AN concentrations were higher in the seasonally frozen soils. Gravel was absent at all sampling sites. Soil particle size analysis (Fig. S2), conducted at 10 cm intervals, revealed low clay content across all samples, with averages of 10.2% in permafrost soils and 6.8% in seasonally frozen soils. Silt and sand content averaged 54.2% and 36.5%, respectively, with a higher proportion of sand in seasonally frozen soils. According to the FAO/USDA soil texture classification (49), sites N1-N4 were classified as silt loam, and site N5 as sandy loam. The topsoil at sites L1, L2, and L3 was classified as sandy loam, with deeper layers identified as silt loam.

### 2.2 Sequence data and diversity of microbial communities

Comprehensive sequencing efforts produced a total of 444,279 high-quality bacterial and archaeal sequences, along with 700,980 micro-eukaryotic sequences, across eight study sites. The average sequence lengths were 453 bp for bacteria and archaea, and 355 bp for micro-eukaryotes. Analysis identified 1,981 bacterial OTUs, 3 archaeal OTUs, and 1,808 eukaryotic OTUs at 97% sequence similarity. Venn diagrams (Fig. 3) revealed that 45.2% (n=896) of bacterial OTUs and 35.0% (n=632) of eukaryotic OTUs were shared between permafrost soils (group N) and seasonally frozen soils (group L). All three archaeal OTUs were shared between N and L groups, with no archaeal OTUs unique to either group, likely due to the low number of archaeal OTUs identified. Upset diagrams (Fig. S3) further highlight the distribution of unique OTUs across the sites, with bacterial unique OTUs ranging from 2 to 18, and micro-eukaryotic unique OTUs ranging from 16 to 79 per site. Notably, the number of endemic OTUs peaked at site N4 for both groups, following a humpbacked pattern correlating with increasing active layer thickness.

**Fig. 3.**
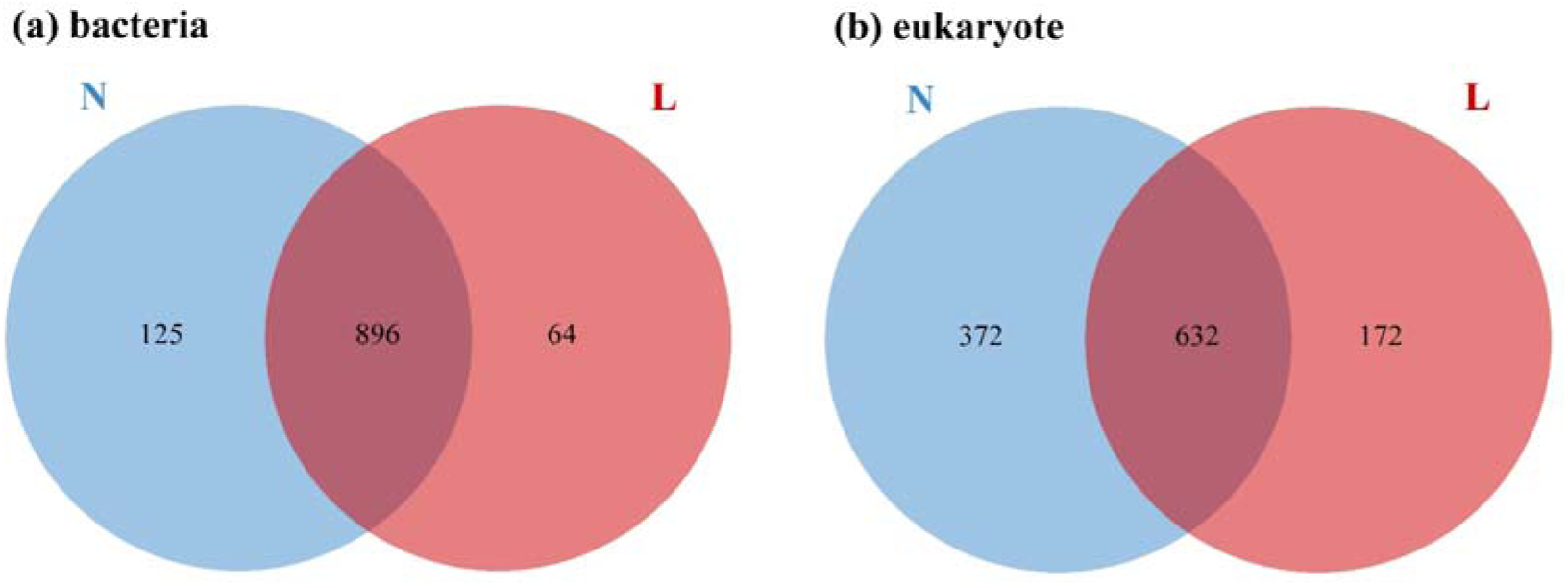
Venn diagram of OTUs among the permafrost soils (N), and seasonally frozen soils (L) for bacteria (a) and micro-eukaryotes (b), with the total number of OTUs in each group shown on the histogram.

Good’s coverage values exceeded 0.99 for all samples (Fig. 4), indicating that the sequencing depth was sufficient to capture microbial diversity. Alpha diversity analysis revealed distinct trends between bacterial and micro-eukaryotic communities in permafrost (N) and seasonally frozen soils (L). In bacterial communities, permafrost soils exhibited higher richness and diversity across all indices compared to seasonally frozen soils (L). Conversely, in micro-eukaryotic communities, seasonally frozen soils displayed slightly higher diversity. Despite these trends, none of the differences between permafrost and seasonally frozen soils were statistically significant.

**Fig. 4.**
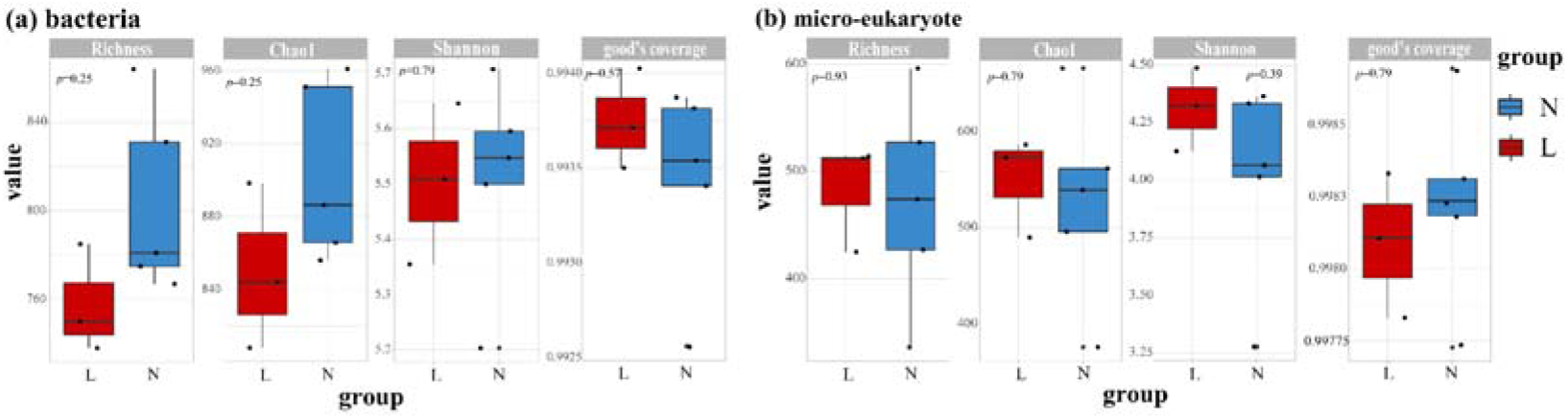
Alpha diversity of permafrost soils (N) and seasonally frozen soils (L) for bacteria (a) and micro-eukaryotes (b).

Non-metric multidimensional scaling (NMDS) plots revealed distinct community structures between bacterial and eukaryotic communities in permafrost versus seasonally frozen soils. Analysis of similarities (ANOSIM) demonstrated significant differences in bacterial community structure across varying permafrost conditions (R = 0.61, *P* < 0.05), whereas no significant differences were observed in the eukaryotic community (R = –0.128, *P* > 0.05) (Fig. 5). These results highlight the contrasting community compositions of soil bacterial and micro-eukaryotic communities in response to permafrost thaw, suggesting that microbial community dynamics vary under different freeze-thaw conditions.

**Fig. 5.**
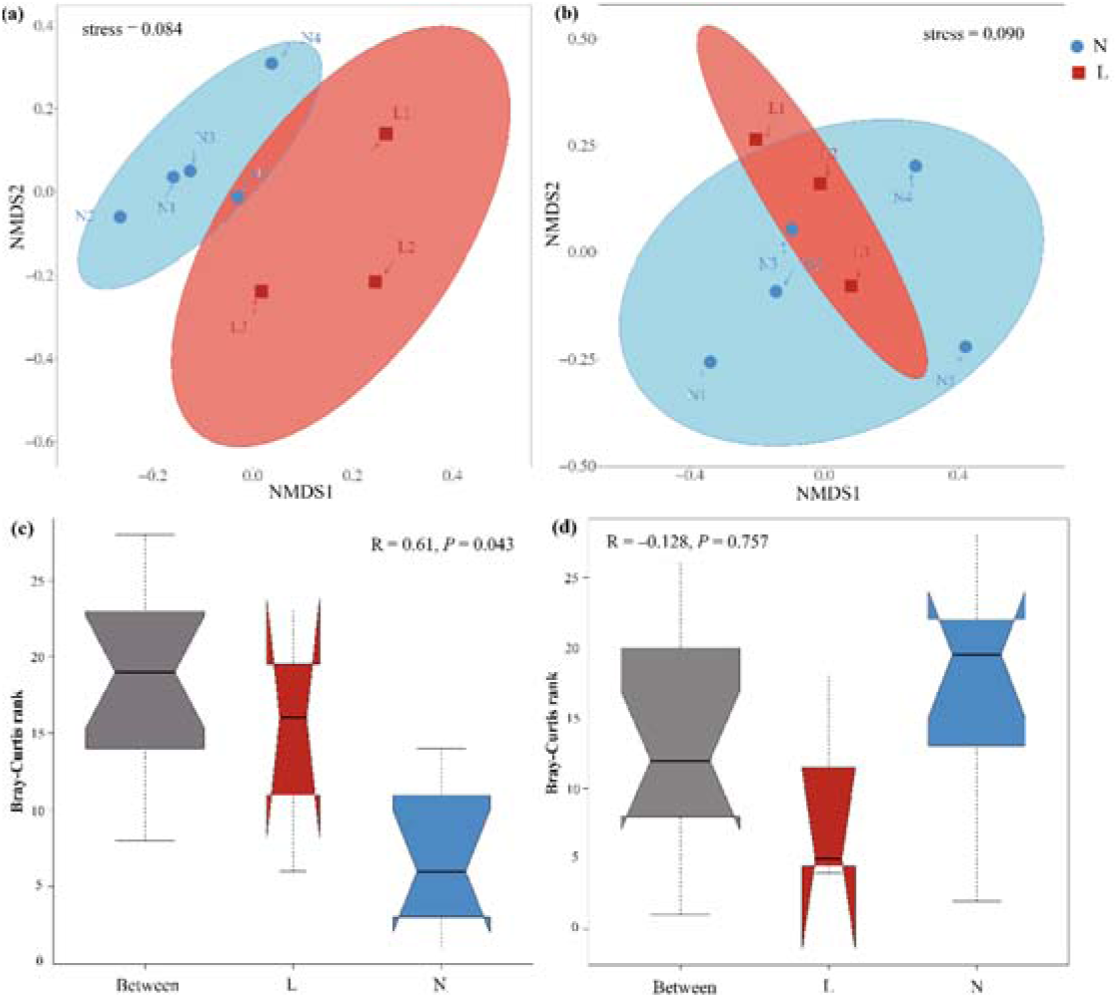
Nonmetric multidimensional scaling (NMDS) plots of bacterial community (a) and eukaryote (b) across the permafrost degradation gradient; Analysis of similarity (ANOSIM) plots of bacterial community (c) and eukaryote (d) across the permafrost degradation gradient. Stress > 0 means

### 2.3 Composition of soil microbial communities

The bacterial component of the soil microbiome was highly diverse, with sequences assigned to 20 phyla, 61 classes, 120 orders, 151 families, and 198 genera. The dominant phyla included Acidobacteria (32.9%), Proteobacteria (32.0%), Actinobacteria (10.0%), Bacteroidetes (9.0%), collectively accounting for 84.0% of all bacterial sequences (Fig. 6(a)). Among these, the phylum Nitrospirae showed a significant decrease along the permafrost degradation gradient (*P* < 0.01). The top four dominant classes accounted for 57.3% of the bacterial community (Fig. 6(b)). Lesser-known phyla made up 75.9% of the total bacterial diversity (Table S2). The top 19 bacterial genera were led by *RB41* (13.0%), *Sphingomonas* (5.3%), *Nitrospira* (4.1%), uncultivated soil bacterium clone C112 (2.3%), and *Terrimonas* (1.1%). Notably, the relative abundance of these genera showed statistically significant differences between permafrost and seasonally frozen soils (Fig. 6(c)), indicating spatial variations in bacterial community compositions that suggest a strong influence of permafrost degradation.

**Fig. 6.**
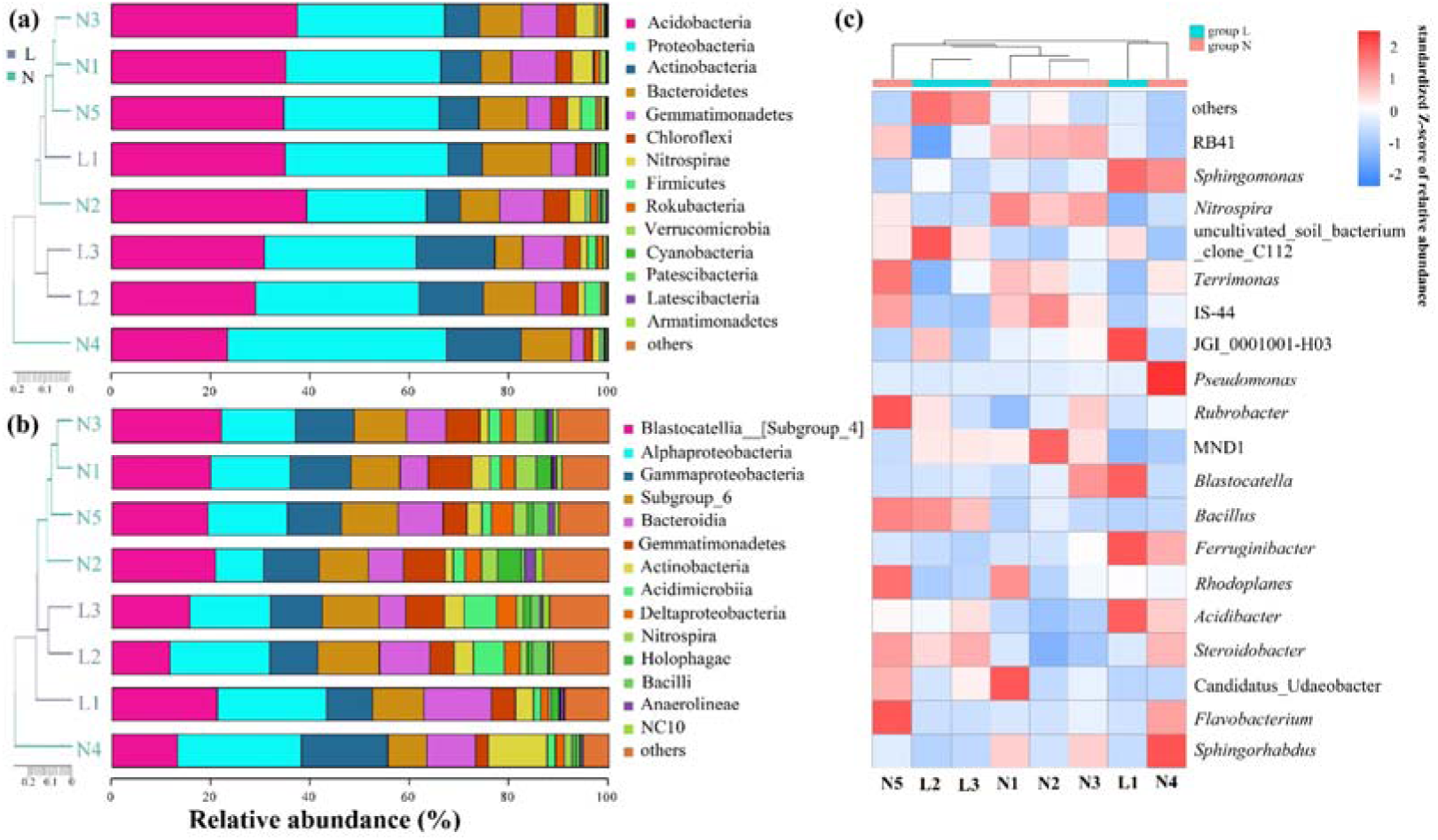
Relative abundance of the soil bacterial community under the permafrost soils (N1-N5) and seasonally frozen soils (L1-L3): (a) Phylum, (b) Class, and (c) Classified Genus. “Others” represents sequences that were unclassified and sequences that were less than 1% of the total.

Archaea were also detected in the soil microbiome, though at a much lower abundance than bacteria. Only three archaeal OTUs were identified (Table S3), all classified within the Thaumarchaeota phylum and Nitrososphaeria class. These OTUs were present in both permafrost and seasonally frozen soils, with none being exclusive to either environment. The low number of archaeal OTUs likely reflects their limited representation in the soil compared to bacteria.

Eukaryotic diversity encompassed six lineages, with Opisthokonta, SAR, and Archaeplastida being the most dominant groups (Fig. 7(a)). Among the 12 eukaryotic phyla identified, Nucletmycea and Rhizaria were predominant, comprising 86.4% of the total eukaryotic sequences. The remaining phyla, including Chloroplastida (8.4%) and Discosea (2.5%), represented a smaller fraction of the community (2.7%) (Fig. 7(b)). Two Phyla, Holozoa and Rhizaria, showed a significant increase in relative abundance with increasing ALT (*P* < 0.05) (Table S4). Heatmap analysis highlighted several dominant fungal families such as Mortierellaceae, Agaricales, and Leptosphaeriaceae, which accounted for 98.1% of the Nucletmycea phylum (Fig. 7(c)). Among the top 20 fungal groups in the micro-eukaryotic community, Hygrophoraceae and Microascaceae showed significant differences between the two soil types (*P* < 0.05) as determined by generalized linear models. Overall, the fungal component of the micro-eukaryotic community showed less variability along the degradation gradient compared to the bacterial community, suggesting a relative stability of fungal populations in response to permafrost changes.

**Fig. 7.**
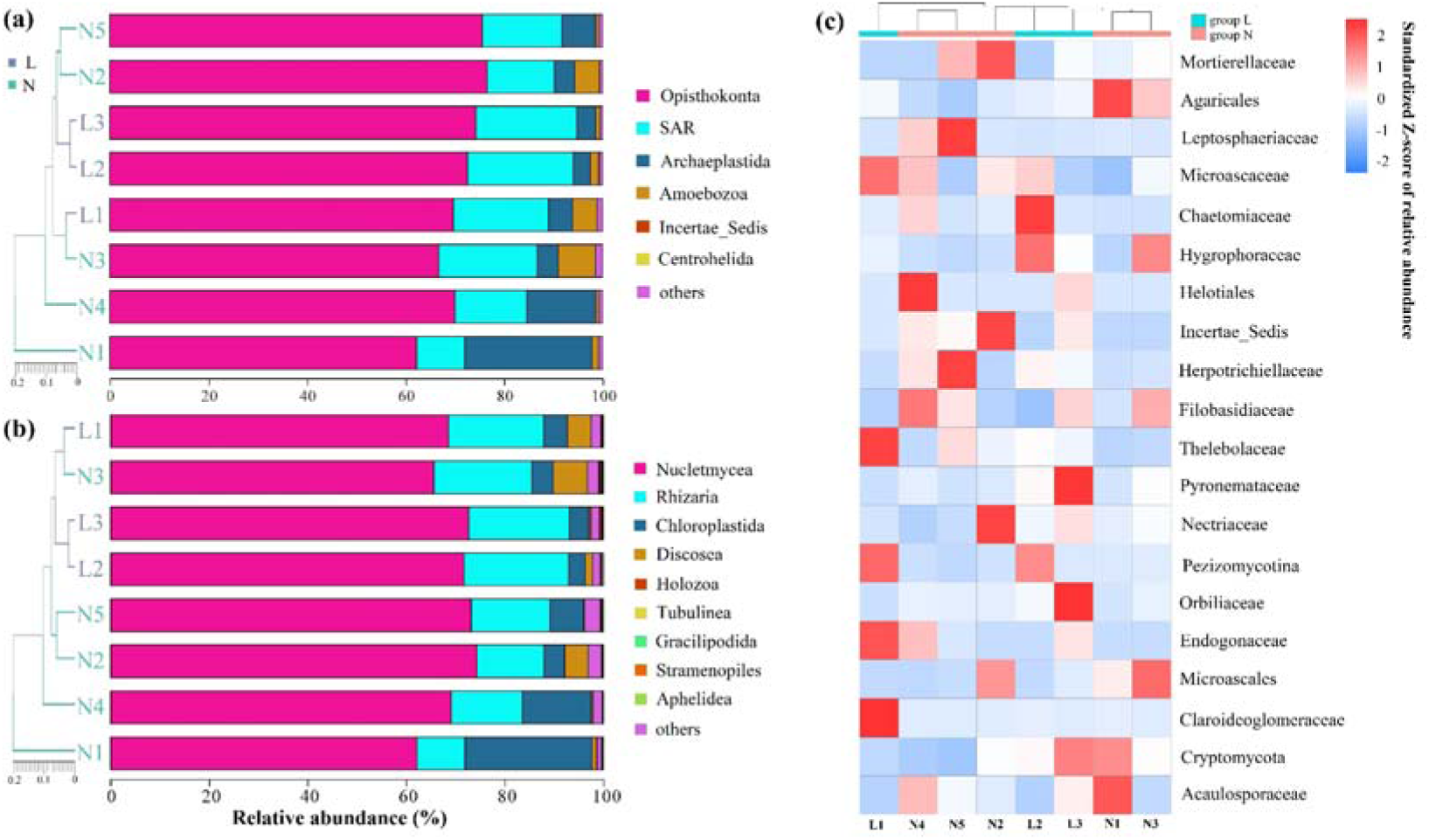
Relative abundance of the eukaryotic community under the permafrost soils (N1–N5) and seasonally frozen soils (L1–L3): (a) Kingdom, (b) Phylum, and (c) Fungal groups of Nucletmycea. “Others” represents sequences that were unclassified and sequences that were less than 1% of the total.

As depicted in Fig. 8(a), LEfSe analysis identified 37 bacterial clades with significant differences across all soil samples, using an LDA threshold of 2.0. In the permafrost soils (N1-N5), 22 bacterial clades were notably enriched, while 15 clades were more abundant in the seasonally frozen soils (L1-L3). Notably, seasonally frozen soils (group L) showed a higher prevalence of members from the Cytophagales order and uncultivated soil bacterium clone C112 at multiple taxonomic levels (order, family, and genus). Conversely, permafrost soils (group N) displayed enrichment in Gammaproteobacteria (class), Nitrosomonadaceae (family), and Nitrospiraceae (family). Previous studies on the Qinghai-Tibet Plateau have identified Cytophagales as potential biomarkers for alpine wetland environments (50).

**Fig. 8.**
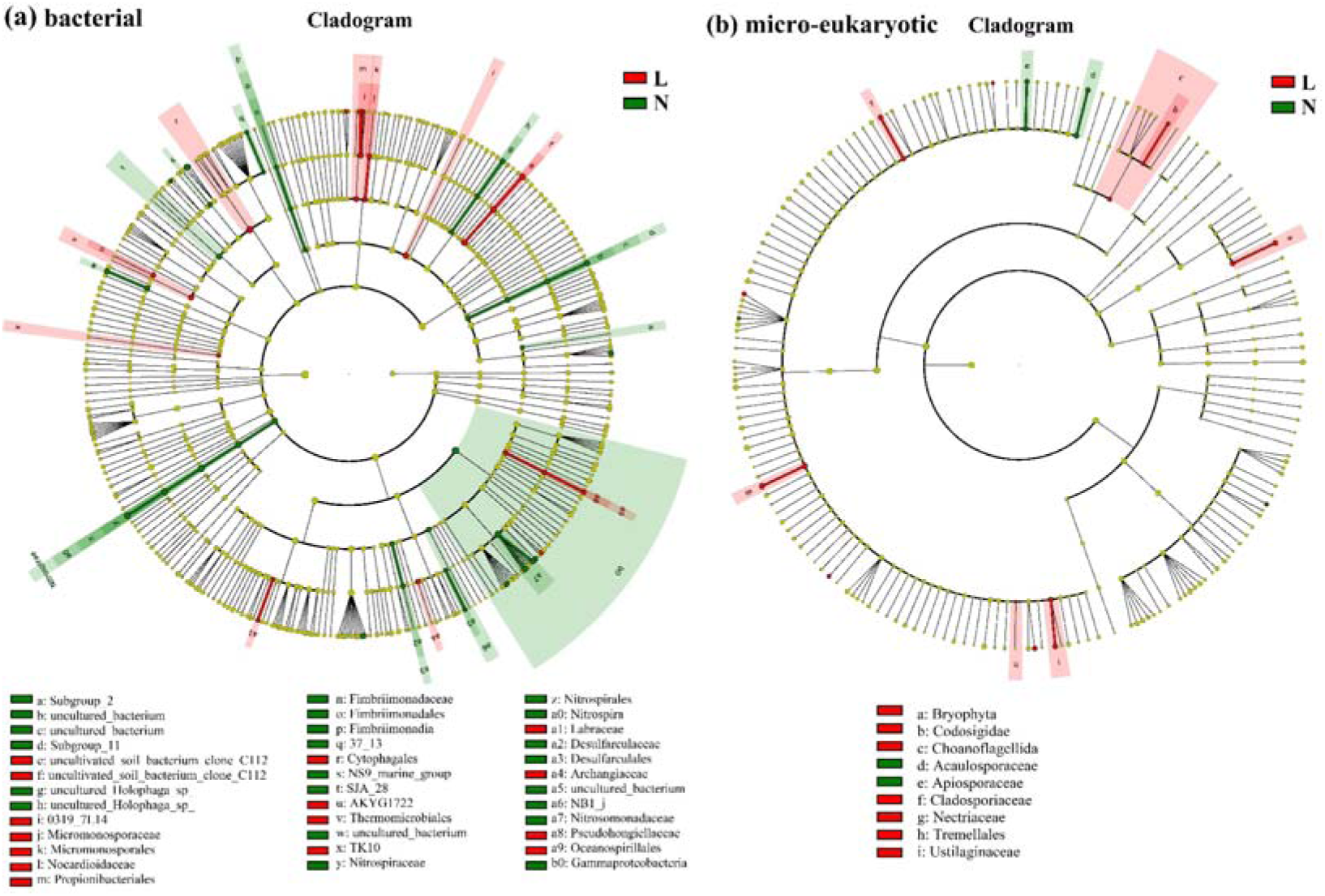
LEfSe analysis and Histogram of LDA scores with a threshold value of 2.0 of soil microbial abundance in permafrost soils (group N) and seasonally frozen soils (group L): soil bacteria (a) and eukaryote (b). Small circles and shading with different colors illustrate the taxa that are enriched in group N and group L.

For the micro-eukaryotic community, as shown in Fig. 8(b), LEfSe analysis revealed six eukaryotic clades—belonging to the Nuclemycea, Rhizaria, Holozoa, and Chloroplastida groups—that were significantly affected by soil degradation. Among these, Acaulosporaceae was identified as a potential biomarker in permafrost soils, with its highest abundance in sample N1 (2.24%). In contrast, Nectriaceae was suggested as a biomarker for seasonally frozen soils, most notably in sample L3 (2.68%). These findings offer insights into the distinct responses of bacterial and micro-eukaryotic communities to environmental changes in permafrost regions, identifying taxa that may serve as indicators of ecological shifts due to permafrost thaw.

### 2.4 Co-occurrence networks in response to permafrost thaw gradient

Co-occurrence networks were constructed for both bacterial and micro-eukaryotic communities to elucidate their interaction patterns in response to the permafrost thaw gradient. These networks were based on robust correlations (R = 0.94) and statistically significant associations (*P* < 0.05) among OTUs, revealing intricate microbial relationships. The bacterial network was notably extensive, comprising 633 nodes connected by 1127 edges, with 54.75% positive and 45.25% negative interactions (Fig. 9(a)). The dominant taxa in the bacterial network, identified by their higher degrees (greater number of connections), included Proteobacteria, Acidobacteria, and Bacteroidetes, which highlighted their central roles in shaping microbial interactions in permafrost soils (51). The node-level topological features of the bacterial and micro-eukaryotic networks were assessed (Table 2), with a modularity index exceeding 0.4, indicating a highly modular network structure with tightly interconnected clusters (52). These modules represented 59.11% of the network, with individual modules containing 4.11% to 8.06% of the total nodes (Fig. 9(b)).

**Fig. 9.**
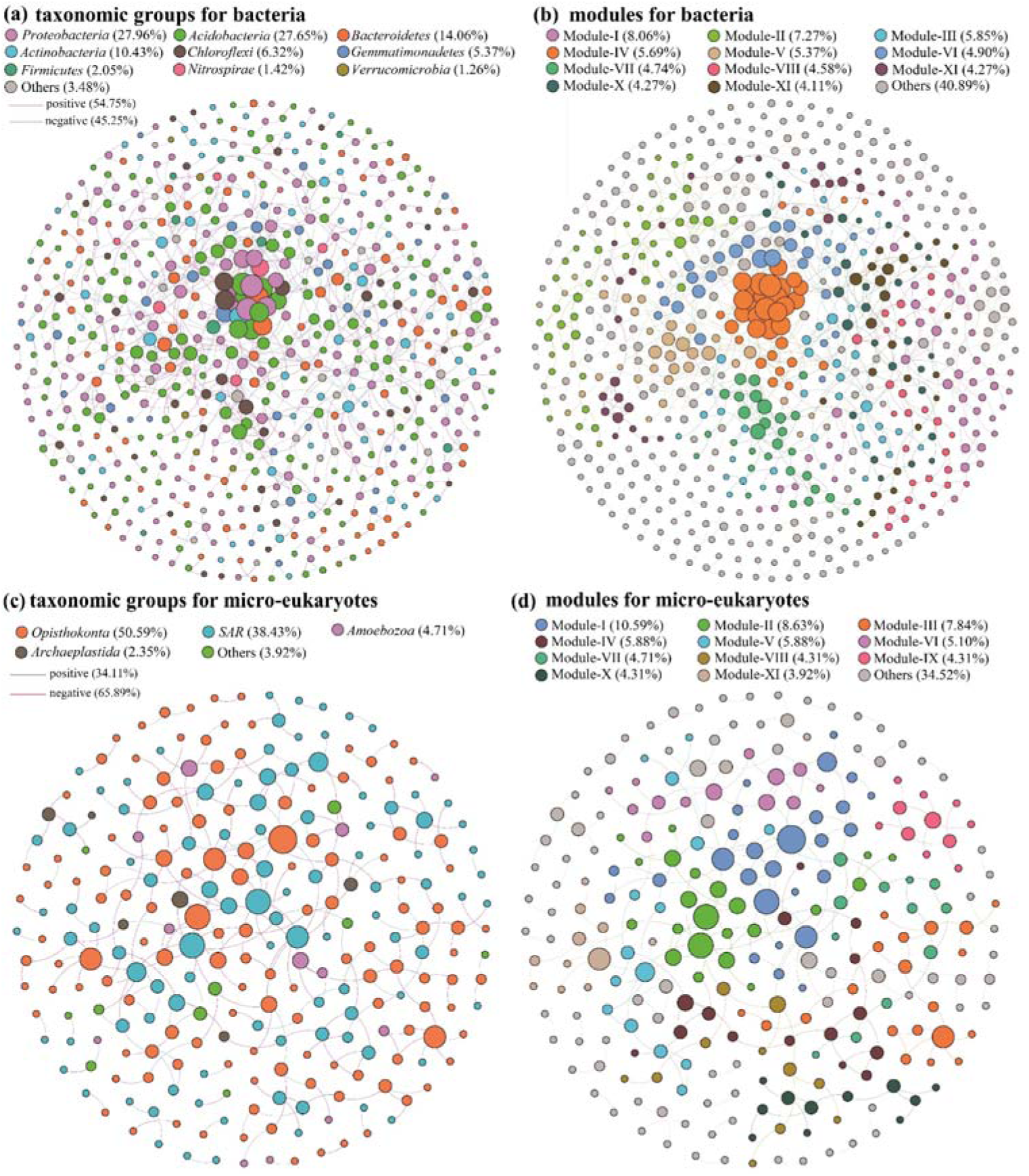
The co-occurrence networks among the OTUs of soil bacterial and micro-eukaryotic communities colored by taxonomic groups ((a), (c)) and modules ((b), (d)) based on Spearman correlation analysis.

**Table 2.** Key topological features of soil microbial networks.

| Topological Parameters | Bacteria | Eukaryota |
| --- | --- | --- |
| Number of edges | 1127 | 258 |
| Number of nodes | 633 | 255 |
| Network density | 0.006 | 0.008 |
| Characteristic degree | 3.561 | 2.024 |
| Characteristic path length | 8.299 | 7.275 |
| Clustering coefficient | 0.369 | 0.125 |
| Modularity | 0.822 | 0.863 |

In contrast, the micro-eukaryotic network, comprising 255 nodes and 258 edges, exhibited a predominance of negative interactions (65.89%) compared to positive ones (34.11%), suggesting more competitive or antagonistic relationships (Fig. 9(c)). The network, primarily consisting of Opisthokonta and SAR taxa, was divided into nine major modules, similar to the bacterial network, which may allow for specialized functional responses to environmental changes (Fig. 9(d)). The contrasting patterns of positive and negative correlations in bacterial and micro-eukaryotic networks underscore the distinct ways these communities respond to environmental stresses associated with permafrost thaw.

### 2.5 Effects of environmental factors on soil microbial communities

The correlation analyses between environmental factors and microbial communities revealed distinct influences on soil bacteria and micro-eukaryotes. Soil chemical properties exhibited mutual positive correlations, whereas elevation, ALT (active layer thickness), soil temperature, and electrical conductivity showed negative relationships with one another. The Mantel test indicated that only a few environmental factors significantly impacted the soil microbial communities (*P* < 0.05). For bacterial communities, soil temperature (r ≥ 0.4, *P* < 0.05) and electrical conductivity (r ≥ 0.4, *P* ≥ 0.05) were the most significant factors, both demonstrating strong correlations. Elevation and vegetation cover (0.2 < r < 0.4, *P* < 0.05) also showed moderate but notable influences. In contrast, the micro-eukaryotic communities were primarily influenced by the carbon-to-nitrogen ratio (r ≥ 0.4, *P* < 0.05) and silt content (0.2 < r < 0.4, *P* < 0.05), suggesting a stronger dependency on soil nutrient dynamics. Interestingly, ALT had a considerable effect on micro-eukaryotes (0.2 < r < 0.4, P ≥ 0.05) but did not significantly influence bacterial communities (Fig. 10(a)).

**Fig. 10.**
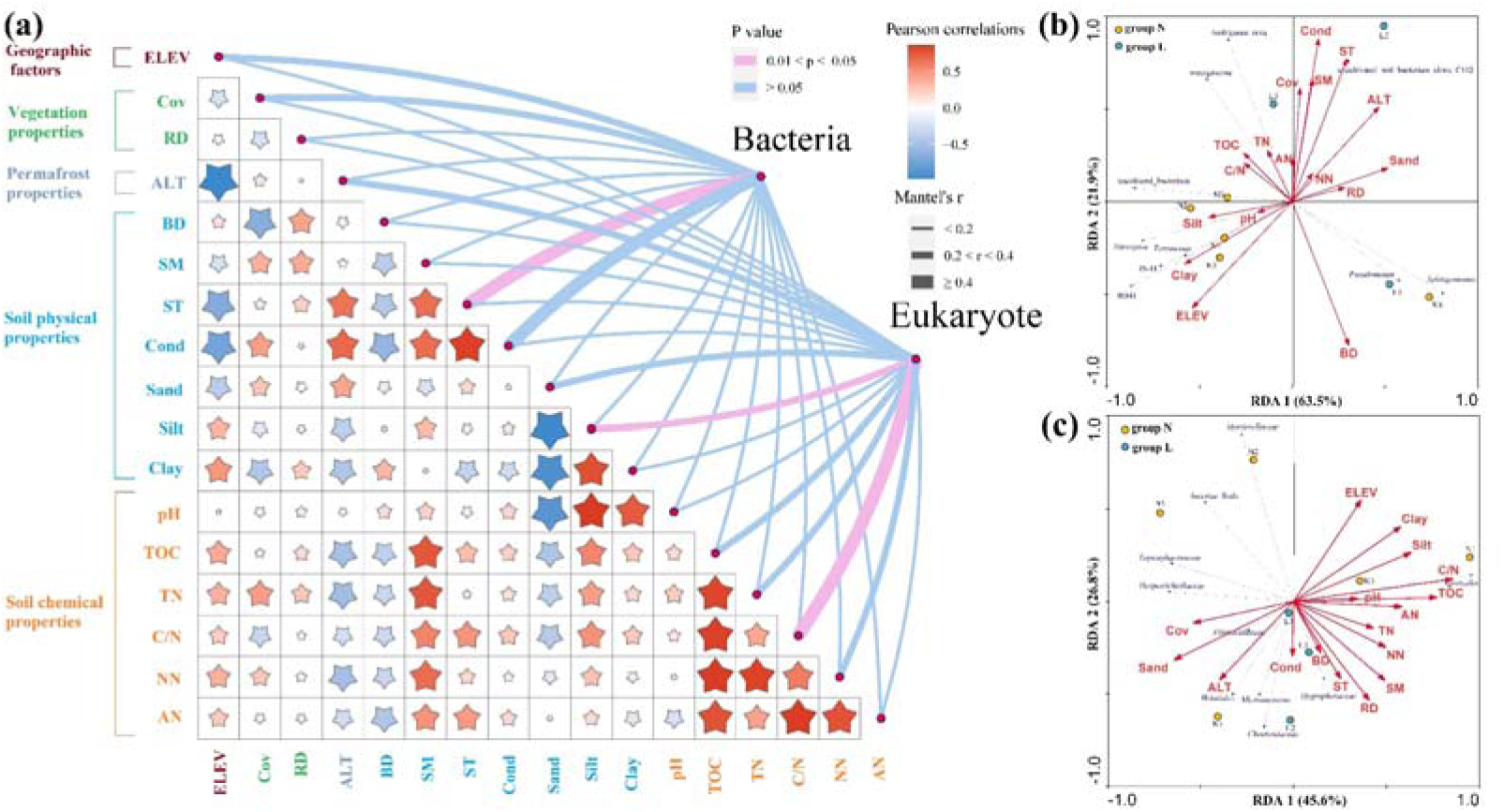
Relationship between environmental factors and soil microbial communities at phylum level revealed by (a) Mantel test and at genus level revealed by RDA analysis for (b) bacterial community and (c) micro-eukaryotic community. ELEV, RD, Cov, ALT, BD, SM, ST, and Cond indicate elevation, root depth, coverage, active layer thickness, bulk density, soil moisture, soil temperature, and electrical conductivity, respectively. TOC, TN, C/N, NN, and AN indicate total organic carbon, total nitrogen, the ratio of C and N, nitrate, and ammonia, respectively. The size of the star is in proportion to the correlation among physicochemical properties. The arrow length corresponds to the variation of microbial community structure that can be explained by the environmental factors and the direction indicates an increasing magnitude of the soil property. Group N and L represent permafrost soils (N1–N5) and seasonally frozen soils (L1**-**L3), respectively.

Redundancy analysis (RDA) further elucidated the relationships between microbial community structure at the genus level and environmental parameters. For bacterial communities, electrical conductivity, soil temperature, and bulk density emerged as the primary driving factors (Fig. 10(b)). Notably, the genus *RB41* (Acidobacteria) exhibited a significant correlation with clay content (*P* < 0.05), suggesting that factors influencing soil aeration and moisture retention may regulate this bacterial genus. Soil texture components, particularly in permafrost soils (group N), played key roles in shaping bacterial community composition, while electrical conductivity was a more influential factor in seasonally frozen soils (group L).

In the micro-eukaryotic community, the carbon-to-nitrogen ratio, root depth, and sand content were the dominant environmental drivers influencing community structure (Fig. 10(c)). These results suggest that soil physical properties had a greater impact on micro-eukaryotic communities than chemical properties, likely due to the more pronounced shifts in soil physical properties along the permafrost thaw gradient.

In summary, our study showed that bacterial communities were more strongly influenced by soil physical properties, while eukaryotic communities were more responsive to variations in soil nutrient content. These findings underscored the distinct ecological niches and adaptive strategies of soil bacterial and micro-eukaryotic communities in permafrost-affected ecosystems.

## 3 Discussion

### 3.1 Impact of permafrost degradation on microbial diversity

Our study highlighted the significant influence of permafrost degradation on bacterial and micro-eukaryotic community diversity in the alpine ecosystems of the Qilian Mountains. While many studies have examined microbial diversity in frozen soil, results have often been inconsistent (53, 54). For instance, research in northern Sweden reported a monotonic decrease in alpha diversity across three stages of permafrost thaw (55), whereas studies from the Qinghai-Tibet plateau observed a contrasting humpback pattern (56). Our findings align with the letter, showing an initial increase followed by a decline in bacterial diversity along the active layer thickness (ALT) gradient. In contrast, micro-eukaryotic diversity exhibited the opposite trend (Fig. 4). This could be attributed to the “mid-dominant effect”, where moderate environmental changes benefit bacterial communities up to a threshold, beyond which diversity declines (57, 58).

Permafrost regions are characterized by harsh conditions, including low temperatures, freeze-thaw cycles, and limited water availability, which support a complex and largely uncharacterized microbial community. In our study, a substantial proportion of soil bacterial (59.1%) and micro-eukaryotic (55.0%) OTUs remained unclassified. The microbial compositions of seasonally frozen and permafrost soils exhibited distinct characteristics (59), with Acidobacteria, Proteobacteria, and Actinobacteria being dominant bacterial phyla (Fig. 6), consistent with other permafrost studies (19, 60). The Actinobacteria/Proteobacteria ratio, often considered an indicator of permafrost degradation, showed an increasing trend with ALT, particularly in seasonally frozen soils. For instance, in sites L2 and L3, this ratio reached 0.39 and 0.52, respectively, compared to 0.26 to 0.34 in permafrost soils (N1-N5). These results align with observations from the Shule River basin (56), indicating that higher Actinobacteria/Proteobacteria ratios may reflect advanced stages of soil degradation and altered microbial community composition in response to thawing. Additionally, *Nitrospira*, a nitrifying bacterial genus, significantly decreased along the permafrost thaw gradient, suggesting a slowdown in nitrogen cycling in these soils (Fig. 6 (b)).

In the micro-eukaryotic community, *Mortierellacea*, a genus involved in chitin decomposition, decreased with increasing ALT (Fig. 7(c)), implying reduced organic matter decomposition and nutrient cycling (61). LEfSe analysis identified GammaProteobateria (Class) and *Nitrospira* (Genus) as bacterial biomarkers in permafrost soils, while Cytophagales (order) dominated seasonally frozen soils. Among micro-eukaryotes, Acaulosporaceae was a biomarker for permafrost soils, and Nectriaceae for seasonally frozen soils (Fig. 8), though regional variations in biomarker detection are common (62).

Our co-occurrence network analysis revealed that bacterial communities maintain a balance of positive and negative interactions, while micro-eukaryotic communities showed more mutual inhibition, indicating differing adaptation strategies within these domains (Fig. 9) (63, 64). This suggests that bacterial communities may be more resilient to environmental disturbances compared to eukaryotic communities, as supported by the relative stability of fungal populations despite temperature fluctuations (47).

Furthermore, our findings suggest that permafrost degradation may destabilize both bacterial and micro-eukaryotic networks. As permafrost degrades, decreased bacterial diversity leads to reduced positive feedback in the network, while increased fungal diversity amplifies negative interactions. A related study in the Qilian Mountains found that network connectivity, centrality, and complexity significantly increased in severely degraded permafrost compared to slightly degraded permafrost (65), implying reduced network stability. These results suggest that permafrost degradation weakens the stability of soil microbial communities.

### 3.2 Responses of microbial communities to environmental changes

The alteration of soil microbial communities in response to permafrost thaw is well established, yet the intricate mechanisms behind these changes remain incompletely understood. Previous research has suggested that warming trends may shift microbial dominance from bacteria to fungi in alpine meadows on the Qinghai-Tibet Plateau (66). Our results reinforced that soil temperature and the carbon-to-nitrogen ratio are critical determinants of microbial community structure (Fig. 10 (a)). Specifically, soil texture and electrical conductivity significantly influenced bacterial communities in permafrost and seasonally frozen soils respectively, while the carbon-to-nitrogen ratio and sand content were key drivers for micro-eukaryotes. To investigate the effects of rising temperature on soil microorganisms in the permafrost regions, a long-term soil experiment conducted in the Alaska tundra found that short-term (1.5 years) temperature increases had little effect on microbial structures, while longer-term warming (4.5 years) significantly impacted these communities (67). In our study, the distribution of soil temperature did not neatly align with ALT gradients, reflecting the complex interplay of elevation and soil conditions. In permafrost soils, soil temperature decreased with lower altitudes, while seasonally frozen soils, it increased due to enhanced soil moisture and electrical conductivity during summer thaw, which likely stimulated microbial metabolism and enzyme activity. Changes in soil water content influence soil permeability and oxygen availability, which can reduce anaerobic microorganisms while promoting aerobic populations, thereby reshaping the composition and structure of microbial communities (47, 69, 70). While nutrient content typically plays a key role in shaping microbial communities, our study revealed that soil physical properties exert a more pronounced influence in this context. Many previous studies were conducted at a basin scale, whereas our sample sites covered an area smaller than 10 km^2^. This smaller geographic scale likely limited the variation in nutrient content, thus amplifying the impact of soil physical characteristics on microbial community response. These findings suggest that microbial responses to environmental factors may vary depending on the spatial scale of the study area.

Although active layer thickness (ALT) was not identified as a dominant factor influencing microbial communities in our study (Fig. 10), changes in soil temperature and carbon-to-nitrogen ratio were directly affected by variations in ALT. Previous research has underscored the critical role of ALT in storing soil organic matter (SOM), which is essential for maintaining soil fertility and supporting ecosystem functions (70, 71). As permafrost degrades, enhanced groundwater flow may promote the mineralization of SOM by improving soil drainage and increasing oxygen availability (72, 73). Consequently, permafrost degradation may reduce the availability of key nutrients such as soil organic carbon (SOC) and total nitrogen (TN), leading to shifts in vegetation and microbial community structures, with potential ecological transitions from highland to drier lowland ecosystems and eventually to wetter communities (74, 75).

These shifts in microbial community composition and diversity, driven by changes in water and nutrient availability due to permafrost degradation, are likely to result in broader ecological changes. Such transformations are often irreversible under current climatic conditions, emphasizing the need for robust ecological security measurements and early warming systems. This study contributes valuable scientific insights into the complex interactions between soil microbial communities and environmental factors in permafrost ecosystems, providing a foundation for further research and policy development aimed at mitigating the impacts of climate change.

## 4 Conclusions

Permafrost plays a crucial role in climatic feedback mechanisms due to its integral influence on water and nutrient cycles. Our study highlights the significant impact of soil properties on the structure and diversity of soil microbial communities within the alpine meadow ecosystem. Specifically, soil temperature was found to be the primary factor shaping bacterial communities, while the carbon-to-nitrogen ratio was more influential in driving micro-eukaryotic community dynamics. The responses of soil bacterial and micro-eukaryotes to permafrost degradation followed distinct patterns: bacterial diversity declined, whereas fungal diversity increased as permafrost thawed. Co-occurrence network analyses revealed that bacterial communities exhibited more positive interactions, suggesting greater stability, while fungal communities demonstrated a higher proportion of negative interactions, indicating a potentially less stable structure. Bacterial networks, with their larger number of nodes and connections, appeared more resilient to environmental changes than their micro-eukaryotic counterparts.

As global warming progresses, microbial community dynamics may shift from bacterial-dominated to eukaryotic-dominated systems. This shift is expected to coincide with reduced stability within the microbial network as soil conditions change. Understanding these shifts is critical for predicting how alpine meadow ecosystems will respond to climate change, providing insights into future ecological transformations and guiding strategies for ecosystem management and preservation in permafrost regions.

## CRediT authorship contribution statement

Zhu Wang: Conceptualization, Methodology, Writing - original draft. Yang Liu: Supervision, Writing - review & editing. Fang Wang: Supervision, Funding acquisition.

## Acknowledgments

The authors gratefully acknowledge to the anonymous referees and the editors for their constructive comments. This work was financially supported by the following foundations: National Key R&D Program (2018YFC0408103); Open Research Fund of the State Key Laboratory of Basin Water Cycle Simulation and Regulation (2016TS01); Major Science and Technology Project of Hunan Province (2018SK1010); and the Industry Science and Technology Plan Project of the Ministry of Water Resources (126301001000160014–2).

## Supplementary materials

Supplementary material associated with this article can be found online at:.

---

**Fig. S1.**
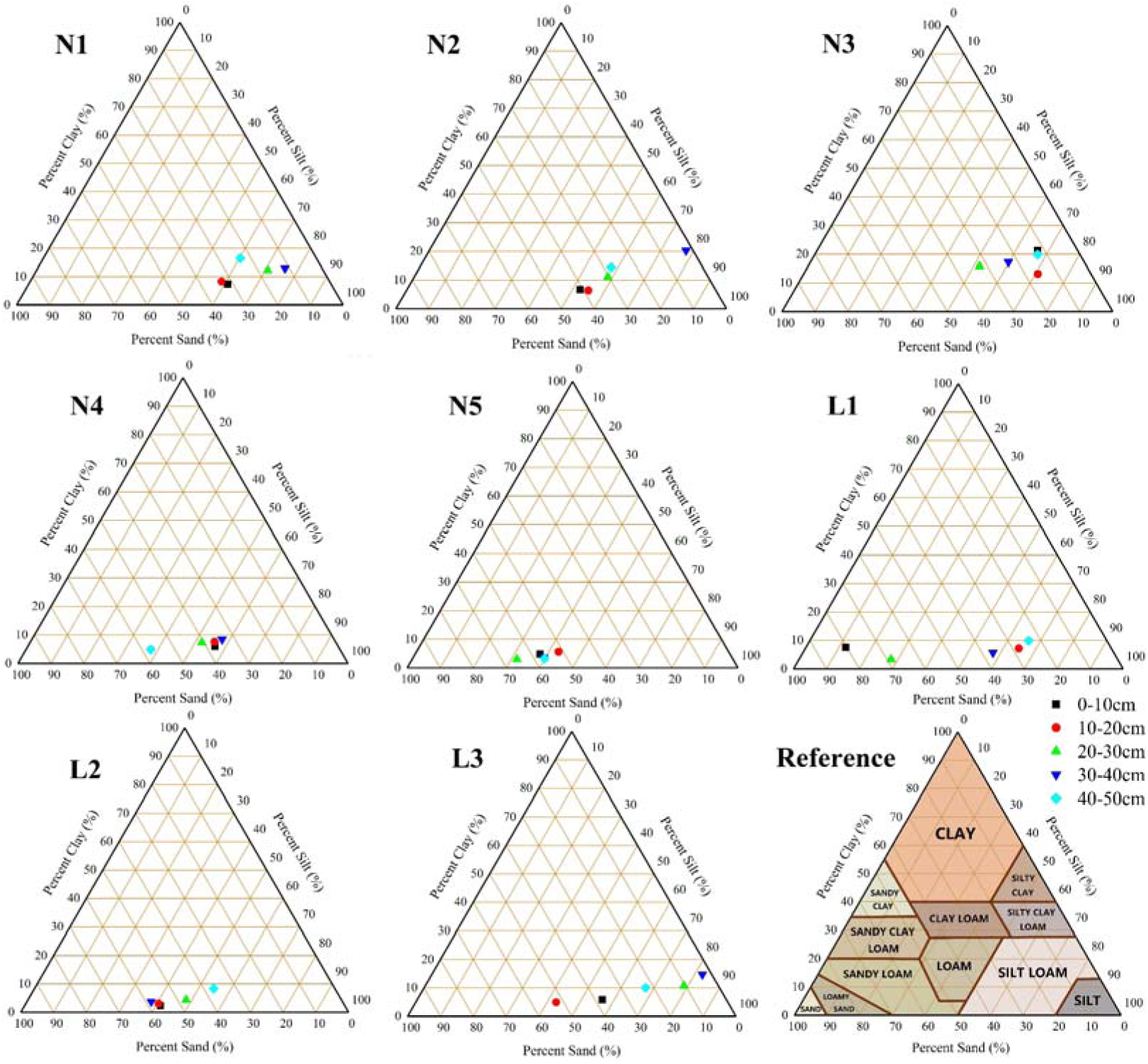
Soil particle size of sampling sites from permafrost soils (N1-N5) and seasonally frozen soils (L1-L3) under different layers (black: 0-10 cm, red: 10-20 cm, green: 20-30 cm, dark blue: 30-40 cm, light blue: 40-50 cm).

**Fig. S2.**
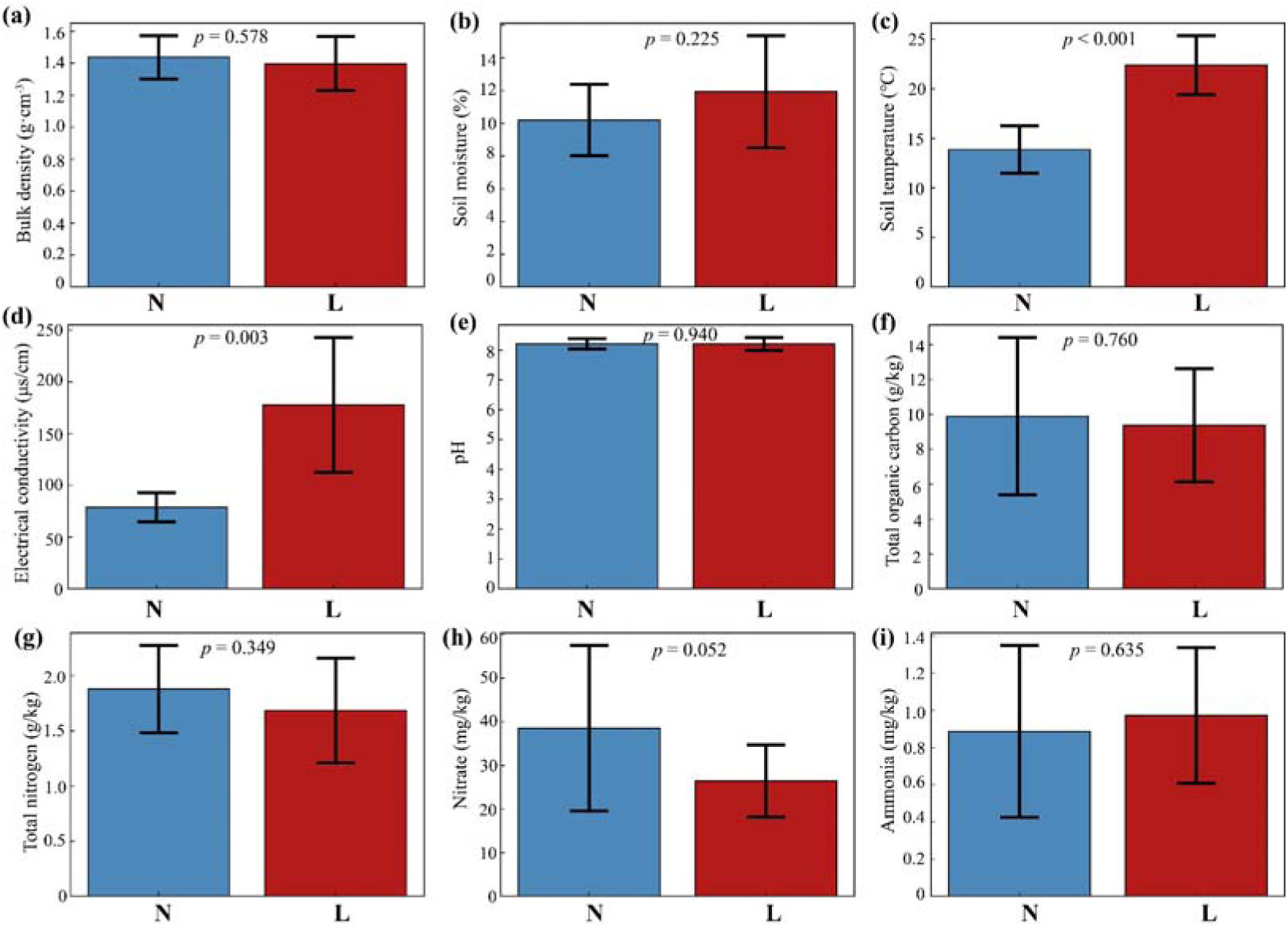
Soil physiochemical properties of sampling sites across permafrost soils (N) and seasonally frozen soils (L) as determined by *t*-test.

**Fig. S3.**
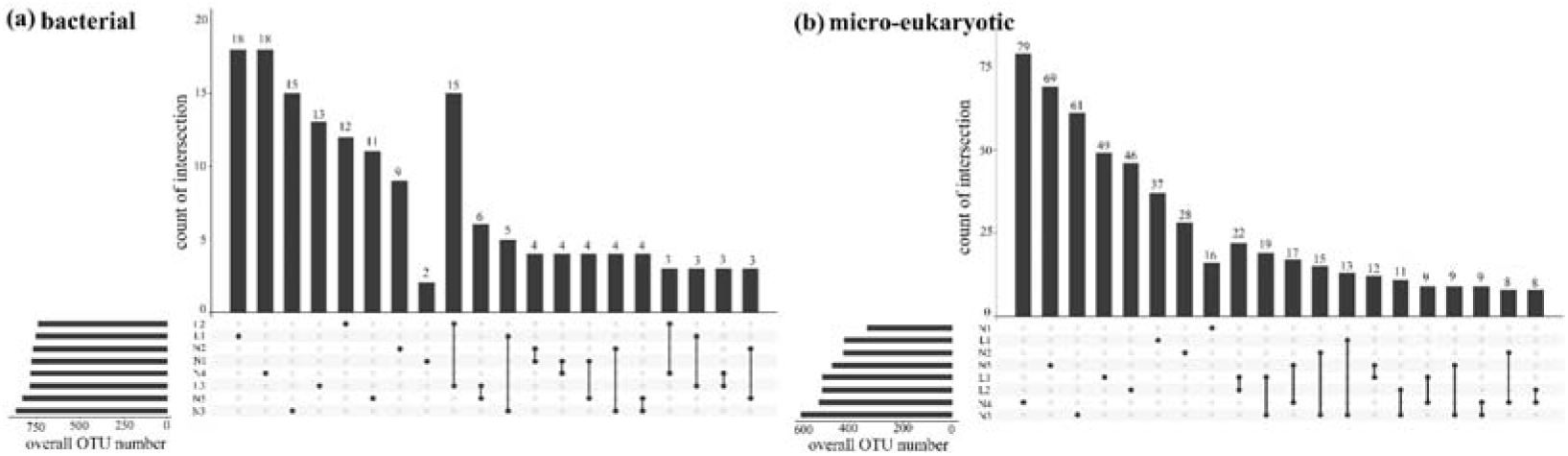
Upset diagrams of OTUs among the permafrost soils (N) and seasonally frozen soils (L) for bacteria (a) and micro-eukaryotes (b).

