## Supplemental Tables 1 to 4, Supplemental Figures 1 to 3 for "Compositional Shifts and Co-occurrence Patterns of Topsoil Bacteria and Micro-Eukaryotes Across a Permafrost Thaw Gradient in Alpine Meadows of the Qilian Mountains, China": 5_Supplementary file.docx

### Table S1. Physical and chemical properties of samples in the permafrost soils (N1-N5) and the seasonally frozen soils (L1-L3).

| Soil  sample | Altitude  (m) | Quadrat  (m) | Coverage  (%) | Dominant  species group | Average root  depth (cm) | ALT  (m) | Soil type | Soil depth  (cm) | BD  (g/cm^3^) | SM  (%) | ST  (℃) | EC  (μs/cm) | pH | TOC  (%) | TN  (mg/kg) | Nitrate  (mg/kg) | Ammonia  (mg/kg) |
| --- | --- | --- | --- | --- | --- | --- | --- | --- | --- | --- | --- | --- | --- | --- | --- | --- | --- |
| N1 | 4103 | 5×5 | 55 | *Polygonum viviparum Rhodiola algida* | 5 | 1.75 | Grass felt soils | 0-20 | 1.27 | 14.6 | 19.7 | 78 | 8.26 | 1.76 | 2379.9 | 60.5 | 2.2 |
|  |  |  |  |  |  |  |  | 20-40 | 1.39 | 13.6 | 16.8 | 94 | 8.19 | 1.60 | 2185.8 | 28.3 | 1.4 |
|  |  |  |  |  |  |  |  | 40-50 | 1.48 | 10.80 | 15.6 | 80 | 8.13 | 1.85 | 2466.3 | 45.1 | 1.2 |
| N2 | 3843 | 5×5 | 65 | *Lagotis brachystachya Leontopodium pusillum* | 2 | 3.46 | Grass felt soils | 0-20 | 1.26 | 10.5 | 16.0 | 91 | 8.22 | 1.08 | 1555.6 | 19.6 | 1.1 |
|  |  |  |  |  |  |  |  | 20-40 | 1.33 | 9.4 | 13.9 | 98 | 8.38 | 0.72 | 2076.6 | 35.4 | 0.9 |
|  |  |  |  |  |  |  |  | 40-50 | 1.34 | 5.0 | 12.1 | 105 | 8.29 | 0.61 | 1001.3 | 10.4 | 0.7 |
| N3 | 3819 | 5×5 | 50 | *Rhodiola algida* | 9 | 3.20 | Grass felt soils | 0-20 | 1.60 | 13.0 | 15.1 | 65 | 8.40 | 1.14 | 2025.8 | 24.8 | 0.6 |
|  |  |  |  |  |  |  |  | 20-40 | 1.58 | 9.6 | 12.9 | 71 | 8.39 | 0.89 | 1984.4 | 19.8 | 0.3 |
|  |  |  |  |  |  |  |  | 40-50 | 1.63 | 9.4 | 11.8 | 74 | 8.32 | 0.69 | 1572.9 | 16.6 | 0.5 |
| N4 | 3766 | 5×5 | 80 | *Potentilla fruticosa* | 5 | 3.20 | Swamp soils | 0-20 | 1.38 | 10.1 | 12.2 | 60 | 8.05 | 1.20 | 1598.7 | 20.5 | 0.6 |
|  |  |  |  |  |  |  |  | 20-40 | 1.54 | 9.9 | 11.5 | 62 | 8.26 | 0.82 | 2167.8 | 39.2 | 0.9 |
|  |  |  |  |  |  |  |  | 40-50 | 1.69 | 10.2 | 9.9 | 59 | 8.45 | 0.35 | 2165.8 | 39.1 | 0.7 |
| N5 | 3767 | 5×5 | 85 | *Potentilla fruticosa* | 3 | 3.53 | Swamp soils | 0-20 | 1.30 | 9.1 | 14.5 | 69 | 7.85 | 1.14 | 1700.8 | 23.1 | 0.7 |
|  |  |  |  |  |  |  |  | 20-40 | 1.34 | 9.6 | 13.5 | 84 | 7.91 | 0.51 | 1998.8 | 32.4 | 1.2 |
|  |  |  |  |  |  |  |  | 40-50 | 1.43 | 8.6 | 12.6 | 91 | 8.07 | 0.47 | 1294.6 | 14.5 | 0.3 |
| L1 | 3496 | 5×5 | 30 | *Artemisia hedinii Elymus dahuricus* | 5 | 4.99 | Dark felty soils | 0-20 | 1.44 | 12.8 | 22.5 | 99 | 8.31 | 0.79 | 1187.9 | 27.7 | 0.8 |
|  |  |  |  |  |  |  |  | 20-40 | 1.67 | 9.3 | 20.0 | 98 | 8.22 | 0.83 | 1054.3 | 15.1 | 0.4 |
|  |  |  |  |  |  |  |  | 40-50 | 1.65 | 4.1 | 16.9 | 98 | 7.80 | 0.20 | 1117.2 | 15.0 | 1.6 |
| L2 | 3403 | 5×5 | 85 | *Carex moorcroftii Plantago depressa* | 8 | 4.94 | Dark felty soils | 0-20 | 1.29 | 11.3 | 25.6 | 154 | 8.08 | 1.21 | 1924.1 | 29.7 | 0.9 |
|  |  |  |  |  |  |  |  | 20-40 | 1.31 | 15.1 | 25.8 | 230 | 8.13 | 1.12 | 2041.0 | 34.0 | 1.4 |
|  |  |  |  |  |  |  |  | 40-50 | 1.25 | 15.2 | 25.8 | 225 | 8.11 | 1.04 | 2047.9 | 34.2 | 1.2 |
| L3 | 3400 | 5×5 | 85 | *Carex moorcroftii* | 3 | 4.95 | Dark felty soils | 0-20 | 1.13 | 11.2 | 24.1 | 188 | 8.13 | 1.36 | 2530.6 | 39.2 | 1.1 |
|  |  |  |  |  |  |  |  | 20-40 | 1.41 | 12.7 | 20.9 | 220 | 8.51 | 0.75 | 1762.4 | 24.7 | 0.8 |
|  |  |  |  |  |  |  |  | 40-50 | 1.43 | 15.7 | 20.0 | 287 | 8.55 | 1.14 | 1508.7 | 18.5 | 0.6 |

^Notes: All values are reported as “mean deviation” based on measurement results for triplicated samples; ALT: active layer thickness; BD - bulk density; SM - soil moisture; ST - soil temperature; EC - electrical conductivity; TOC - total organic carbon; TN - total nitrogen.^

### Table S2. Relative abundance of bacterial community at the phylum level.

| OTUID | N1 | N2 | N3 | N4 | N5 | L1 | L2 | L3 |
| --- | --- | --- | --- | --- | --- | --- | --- | --- |
| Acidobacteria | 35.0123 | 38.9194 | 37.1672 | 23.2544 | 34.5991 | 34.8606 | 28.8927 | 30.6501 |
| Proteobacteria | 31.1418 | 23.9395 | 29.5988 | 44.0347 | 31.2464 | 32.7109 | 32.9411 | 30.5560 |
| Actinobacteria | 8.0130 | 6.7054 | 6.9460 | 14.9537 | 7.9450 | 6.7838 | 12.8459 | 15.7906 |
| Bacteroidetes | 6.1509 | 7.8613 | 8.4419 | 10.1051 | 9.6396 | 13.8972 | 10.5079 | 5.6070 |
| Gemmatimonadetes | 8.9544 | 8.8028 | 7.0663 | 2.5629 | 4.7910 | 4.9427 | 5.3350 | 8.3059 |
| Chloroflexi | 3.2010 | 5.0839 | 3.7659 | 1.6423 | 3.2952 | 3.1592 | 3.2010 | 3.2115 |
| Nitrospirae | 4.2523 | 3.2690 | 3.8130 | 1.5377 | 2.7250 | 0.5597 | 1.3913 | 1.4593 |
| Firmicutes | 0.2040 | 0.8526 | 0.4603 | 0.7584 | 3.1016 | 0.2615 | 3.1330 | 1.8045 |
| Rokubacteria | 1.1298 | 1.5744 | 0.9310 | 0.2824 | 0.8787 | 0.1308 | 0.7479 | 1.3704 |
| Verrucomicrobia | 1.2919 | 0.4812 | 0.5753 | 0.3347 | 0.9990 | 0.4498 | 0.4184 | 0.6590 |
| Cyanobacteria | 0 | 0.3452 | 0.1831 | 0.0052 | 0.0157 | 1.6319 | 0.0209 | 0.0366 |
| Patescibacteria | 0.1517 | 0.7898 | 0.1935 | 0.2301 | 0.1569 | 0.1569 | 0.2929 | 0.2354 |
| Thaumarchaeota | 0.1674 | 0.8264 | 0.3766 | 0.1203 | 0.1778 | 0.2040 | 0.1308 | 0.1883 |
| Latescibacteria | 0.1831 | 0.3923 | 0.2615 | 0.0575 | 0.2824 | 0.0366 | 0.0366 | 0.0941 |
| Armatimonadetes | 0.0471 | 0.1046 | 0.0732 | 0.0209 | 0.0366 | 0.0732 | 0 | 0.0105 |
| WS2 | 0.0262 | 0.0209 | 0.0366 | 0.0418 | 0.0366 | 0.0575 | 0.0732 | 0.0209 |
| norank | 0.0418 | 0.0209 | 0.0628 | 0.0314 | 0.0052 | 0.0575 | 0.0262 | 0 |
| Entotheonellaeota | 0.0262 | 0.0052 | 0.0314 | 0.0052 | 0.0575 | 0.0157 | 0 | 0 |
| Fibrobacteres | 0 | 0 | 0.0105 | 0.0052 | 0 | 0.0105 | 0 | 0 |
| Elusimicrobia | 0.0052 | 0 | 0.0052 | 0 | 0.0052 | 0 | 0.0052 | 0 |
| Thermotogae | 0 | 0 | 0 | 0.0157 | 0 | 0 | 0 | 0 |
| BRC1 | 0 | 0.0052 | 0 | 0 | 0.0052 | 0 | 0 | 0 |

### Table S3. Relative abundance of archaeal community at OTU level.

|  | N1 | N2 | N3 | N4 | N5 | L1 | L2 | L3 |
| --- | --- | --- | --- | --- | --- | --- | --- | --- |
| OTU86 | 0.8750 | 0.9620 | 0.8750 | 0.9565 | 0.7353 | 0.8205 | 0.7200 | 0.6389 |
| OTU225 | 0.1250 | 0.0253 | 0.1111 | 0.0435 | 0.1765 | 0.1026 | 0.2400 | 0.3056 |
| OTU740 | 0.0000 | 0.0127 | 0.0139 | 0.0000 | 0.0882 | 0.0769 | 0.0400 | 0.0556 |

### Table S4. Relative abundance of eukaryotic community at phylum level.

|  | N1 | N2 | N3 | N4 | N5 | L1 | L2 | L3 |
| --- | --- | --- | --- | --- | --- | --- | --- | --- |
| Nucletmycea | 62.0721 | 74.2592 | 65.5155 | 69.0278 | 73.0970 | 68.5118 | 71.6435 | 72.6218 |
| Rhizaria | 9.7962 | 13.6124 | 19.8784 | 14.4988 | 15.9778 | 19.2986 | 21.2067 | 20.4302 |
| Chloroplastida | 25.7484 | 4.1918 | 4.3067 | 13.9675 | 6.8126 | 4.8253 | 3.3999 | 3.8648 |
| Discosea | 0.9477 | 4.7844 | 6.9454 | 0.3551 | 0.2146 | 4.7972 | 1.4918 | 0.4470 |
| No_rank | 0.9094 | 2.6847 | 2.2888 | 1.7906 | 3.1777 | 1.9720 | 1.3487 | 1.7957 |
| Holozoa | 0.0485 | 0.2069 | 0.2938 | 0.0562 | 0.2503 | 0.2529 | 0.2733 | 0.4419 |
| Tubulinea | 0.1584 | 0.1482 | 0.3065 | 0.1073 | 0.0588 | 0.1660 | 0.2197 | 0.1379 |
| Gracilipodida | 0.2197 | 0.0536 | 0.2299 | 0.0996 | 0.3116 | 0.0077 | 0.0562 | 0.1890 |
| Stramenopiles | 0.0332 | 0.0179 | 0.0511 | 0.0230 | 0.0588 | 0.0307 | 0.0868 | 0.0409 |
| Aphelidea | 0.0307 | 0.0102 | 0.1073 | 0.0255 | 0 | 0.0741 | 0.0383 | 0.0051 |
| uncultured | 0.0204 | 0.0255 | 0.0434 | 0.0460 | 0.0332 | 0.0485 | 0.0102 | 0.0230 |
| Alveolata | 0 | 0 | 0 | 0 | 0 | 0 | 0.2197 | 0.0026 |
| Apusomonadidae | 0.0153 | 0.0051 | 0.0332 | 0.0026 | 0.0077 | 0.0153 | 0.0051 | 0 |

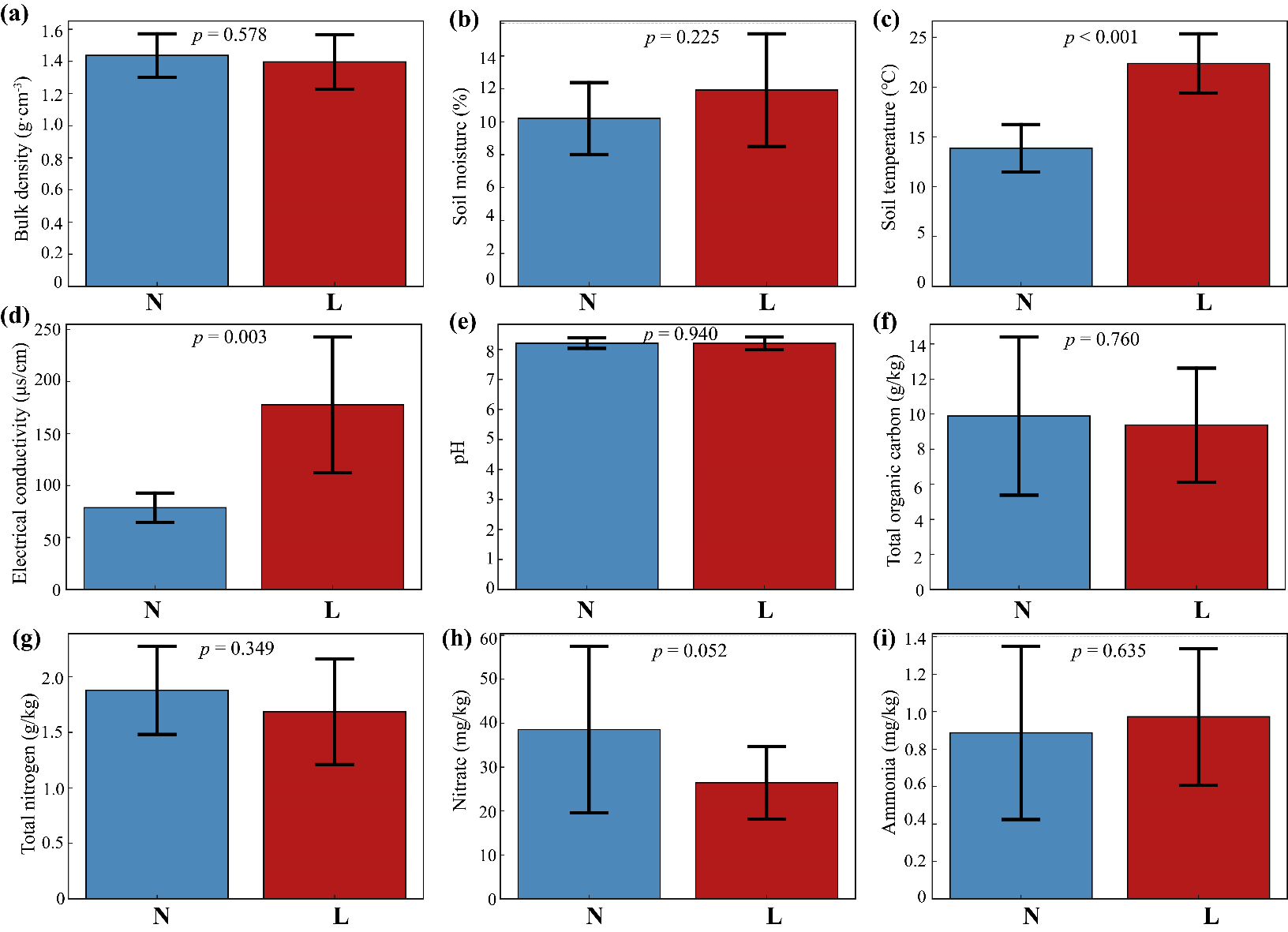

### Fig. S1. Soil physiochemical properties of sampling sites across permafrost soils (N) and seasonally frozen soils (L) as determined by *t*-test.

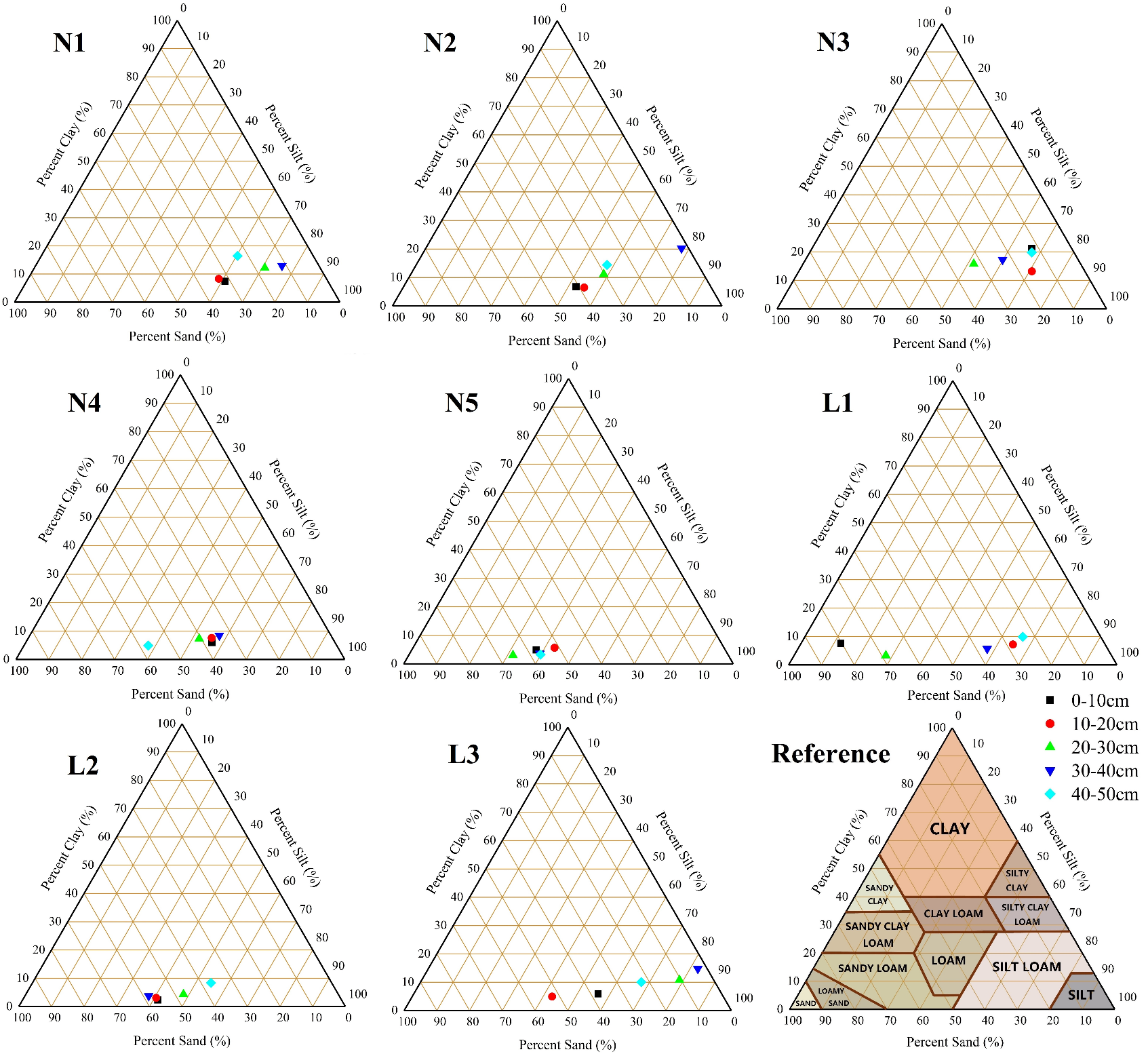

### Fig. S2. Soil particle size of sampling sites from permafrost soils (N1-N5) and seasonally frozen soils (L1-L3) under different layers (black: 0-10 cm, red: 10-20 cm, green: 20-30 cm, dark blue: 30-40 cm, light blue: 40-50 cm).

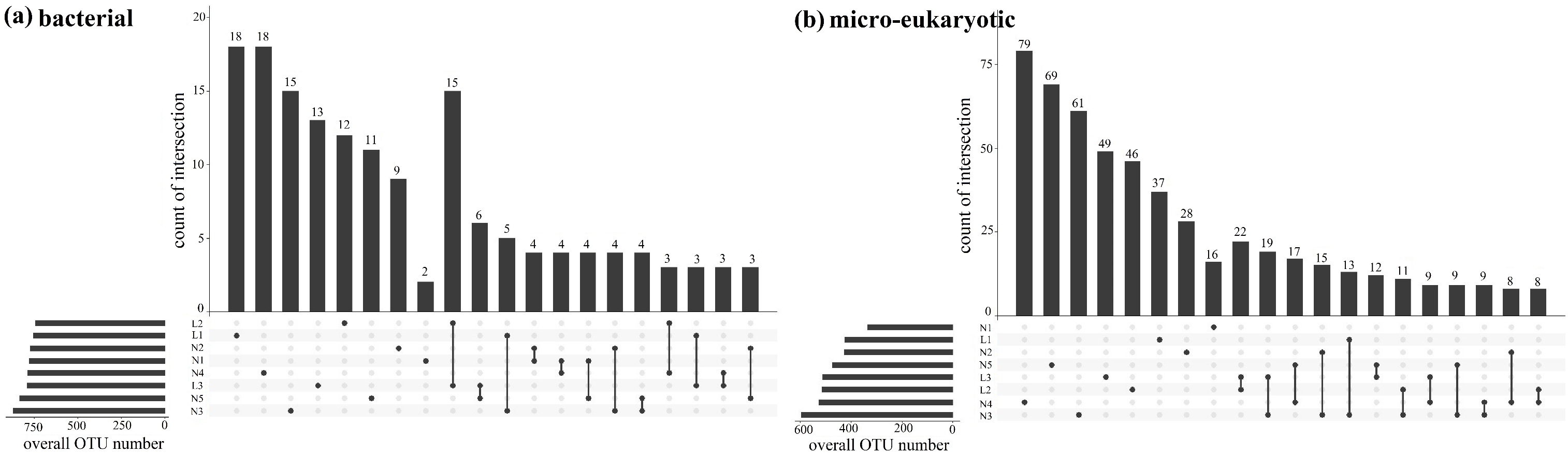

### Fig. S3. Upset diagrams of OTUs among the permafrost soils (N) and seasonally frozen soils (L) for bacteria (a) and micro-eukaryotes (b).
